# Enhancer AAV toolbox for accessing and perturbing striatal cell types and circuits

**DOI:** 10.1101/2024.09.27.615553

**Authors:** Avery C. Hunker, Morgan E. Wirthlin, Gursajan Gill, Nelson J. Johansen, Marcus Hooper, Victoria Omstead, Sara Vargas, M. Nathaly Lerma, Naz Taskin, Natalie Weed, William D. Laird, Yemeserach M. Bishaw, Jacqueline L. Bendrick, Bryan B. Gore, Yoav Ben-Simon, Ximena Opitz-Araya, Refugio A. Martinez, Sharon W. Way, Bargavi Thyagarajan, Sven Otto, Raymond E.A. Sanchez, Jason R. Alexander, Avalon Amaya, Adam Amster, Joel Arbuckle, Angela Ayala, Pam M. Baker, Tyler Barcelli, Stuard Barta, Darren Bertagnolli, Cameron Bielstein, Prajal Bishwakarma, Jessica Bowlus, Gabriella Boyer, Krissy Brouner, Brittny Casian, Tamara Casper, Anish Bhaswanth Chakka, Rushil Chakrabarty, Michael Clark, Kaity Colbert, Scott Daniel, Tim Dawe, Maxwell Departee, Peter DiValentin, Nicholas P. Donadio, Nadezhda I. Dotson, Deepanjali Dwivedi, Tom Egdorf, Tim Fliss, Amanda Gary, Jeff Goldy, Conor Grasso, Erin L. Groce, Kathryn Gudsnuk, Warren Han, Zeb Haradon, Sam Hastings, Olivia Helback, Windy V. Ho, Cindy Huang, Tye Johnson, Danielle L. Jones, Zoe Juneau, Jaimie Kenney, Madison Leibly, Su Li, Elizabeth Liang, Henry Loeffler, Nicholas A. Lusk, Zachary Madigan, Jessica Malloy, Jocelin Malone, Rachel McCue, Jose Melchor, John K. Mich, Skyler Moosman, Elyse Morin, Robyn Naidoo, Dakota Newman, Kiet Ngo, Katrina Nguyen, Aaron L. Oster, Ben Ouellette, Alana A. Oyama, Nick Pena, Trangthanh Pham, Elliot Phillips, Christina Pom, Lydia Potekhina, Shea Ransford, Melissa Reding, Dean F. Rette, Cade Reynoldson, Christine Rimorin, Ana Rios Sigler, Dana B. Rocha, Kara Ronellenfitch, Augustin Ruiz, Lane Sawyer, Josh Sevigny, Nadiya V. Shapovalova, Noah Shepard, Lyudmila Shulga, Sherif Soliman, Brian Staats, Michael J. Taormina, Michael Tieu, Yimin Wang, Josh Wilkes, Toren Wood, Thomas Zhou, Ali Williford, Nick Dee, Tyler Mollenkopf, Lydia Ng, Luke Esposito, Brian Kalmbach, Shenqin Yao, Jeanelle Ariza, Forrest Collman, Shoaib Mufti, Kimberly Smith, Jack Waters, Ina Ersing, Marcella Patrick, Hongkui Zeng, Ed S. Lein, Yoshiko Kojima, Greg Horwitz, Scott F. Owen, Boaz P. Levi, Tanya L. Daigle, Bosiljka Tasic, Trygve E. Bakken, Jonathan T. Ting

**Affiliations:** Allen Institute for Brain Science, Seattle, WA; Department of Neurosurgery, Stanford University School of Medicine, Stanford, CA; Allen Institute for Neural Dynamics, Seattle, WA; Department of Neurobiology & Biophysics, University of Washington, Seattle, WA; Addgene Watertown, MA; Department of Neurological Surgery, University of Washington, Seattle, WA; Department of Otolaryngology, Head and Neck Surgery, University of Washington, Seattle, WA; Washington National Primate Research Center, Seattle, WA

**Keywords:** Medium spiny neuron, spiny projection neuron, cholinergic, Sst-chodl, Pvalb, basal ganglia, caudoputamen, Adeno-associated virus, Addgene, striatonigral, striatopallidal, macaque

## Abstract

We present an enhancer AAV toolbox for accessing and perturbing striatal cell types and circuits. Best-in-class vectors were curated for accessing major striatal neuron populations including medium spiny neurons (MSNs), direct and indirect pathway MSNs, as well as Sst-Chodl, Pvalb-Pthlh, and cholinergic interneurons. Specificity was evaluated by multiple modes of molecular validation, three different routes of virus delivery, and with diverse transgene cargos. Importantly, we provide detailed information necessary to achieve reliable cell type specific labeling under different experimental contexts. We demonstrate direct pathway circuit-selective optogenetic perturbation of behavior and multiplex labeling of striatal interneuron types for targeted analysis of cellular features. Lastly, we show conserved *in vivo* activity for exemplary MSN enhancers in rat and macaque. This collection of striatal enhancer AAVs offers greater versatility compared to available transgenic lines and can readily be applied for cell type and circuit studies in diverse mammalian species beyond the mouse model.

## INTRODUCTION

The striatum, a key component of the basal ganglia, plays a critical role in motor control, habitual behaviors, and cognitive functions, and its dysfunction is central to a variety of movement disorders and substance use disorders^1^ ^4^. The striatum is comprised of diverse neuronal populations, including the highly abundant medium spiny neurons (MSNs) and relatively rare local interneurons, each contributing important roles within functionally distinct circuits^5^ ^8^. MSNs are the principal GABAergic projection neurons of the striatum and are divided roughly equally between Drd1 and Drd2 receptor expressing neuron subtypes (D1 MSNs and D2 MSNs, respectively). Effective targeting and manipulation of these distinct cell types is crucial for unraveling their specific contributions to functional circuits and for developing effective therapeutic interventions to treat basal ganglia disorders.

Considerable progress has been made with respect to delineating cellular and synaptic properties and dissecting the functional role of distinct striatal cell types to mouse behavior. This rapid progress has largely been attributable to the development of transgenic mouse lines, used alone or in combination with viral labeling strategies, to achieve reliable genetic targeting and perturbation of striatal circuit components. For example, cell type-specific bacterial artificial chromosome (BAC) transgenic GFP reporter and Cre driver lines from the <u>G</u>ene <u>E</u>xpression <u>N</u>ervous <u>S</u>ystem <u>AT</u>las (GENSAT) project (www.gensat.org)^9^ provided a means by which researchers could directly test foundational circuit models about the role of the canonical direct (striatonigral) versus indirect (striatopallidal) basal ganglia pathways in motor initiation, coordination, and dysfunction. Drd1-GFP and Drd1-Cre BAC transgenic lines were initially developed for targeting direct pathway D1 MSNs, and Drd2-GFP and Drd2-Cre lines for targeting indirect pathway D2 MSNs^10,11^. Many other derivative BAC transgenic mouse lines followed, offering improved specificity of labeling (e.g., Adora2a-Cre^9^) or alternate transgene cargos (e.g., D1 and D2 BAC trap lines^12^ or Drd1a-tdTomato line^13^) for diversifying experimental applications. Similarly, various other transgenic mouse lines have been utilized to target the diverse interneuron types of the striatum, often with cell type discrimination based on a combination of distinctive neurochemical, morphological, and electrophysiological signatures^6,14,15^.

Importantly, none of the foundational transgenic mouse lines mentioned above are applicable for research in other species. AAV vectors paired with compact cell type-specific enhancers can offer a promising alternative to mouse transgenic technologies to bridge this important gap and to enable genetic targeting of homologous cell types both in the mouse and across additional mammalian model species. Notably, enhancer AAVs can be used in wild-type animals and do not require complicated breeding of transgenic mouse lines. We and others have demonstrated feasibility of enhancer AAVs for targeting diverse cortical neuron types in mice, rats, ferrets, marmosets, macaques, and human *ex vivo* brain slices derived from neurosurgeries^16^ ^26^. Notably, cell type specific enhancer discovery is directly enabled by emerging large-scale single cell epigenetic datasets, especially single cell (sc) and single nucleus (sn) <u>A</u>ssay for <u>T</u>ransposase <u>A</u>ccessible <u>C</u>hromatin with sequencing (sc/snATAC-seq), aligned to brain cell type taxonomies from sc/snRNA-seq. Foundational striatal cell type taxonomies are currently available to compare and align for the mouse^27^ ^30^, marmoset^31^, macaque^32^, and human^33,34^, and serve as a foundational transcriptomic framework for future viral genetic tool molecular validation and use for biological discovery.

Our study aimed to develop and validate an enhancer AAV toolbox for targeting several major striatal neuron populations, including pan-MSNs, direct and indirect pathway MSNs, and Sst-Chodl, Pvalb-Pthlh, and cholinergic interneurons. Using enhancer elements derived from bulk and single-cell ATAC-seq data, we engineered AAV vectors that demonstrate high specificity in labeling these neuronal populations in the mouse brain, with molecular validation anchored in the Allen Institute mouse whole brain hierarchical cell type taxonomy at subclass resolution. We evaluated these vectors for their strength and specificity through various routes of administration and functional assays, including the use of Cre and Flp recombinases, optogenetic actuators, and genetically encoded calcium indicators. Moreover, we investigated suitability of these enhancer AAV vectors for cross-species applications by comparing *in vivo* chromatin accessibility for select MSN enhancers across mouse, rat, macaque, and human, and conducting *in vivo* characterization of enhancer AAV activity in rat and macaque brain. The high conservation of enhancer activity across species underscores the immense potential of these tools for new directions in comparative cellular neuroscience and translational research.

Enhancer AAV screening and validation data from this collection can be viewed in the Allen Institute’s Genetic Tools Atlas (GTA) at https://portal.brain-map.org/genetic-tools/genetic-tools-atlas, an open access community resource created with the joint support of the NIH BRAIN Armamentarium consortium.

## RESULTS

### Identification and *in vivo* validation of striatal subclass specific enhancers from diverse epigenetic datasets

To create a striatum-centric enhancer AAV toolbox, we adopted a one-at-a-time screening pipeline evaluating putative enhancer sequences for driving brain region and cell type-specific SYFP2 reporter expression (**Figure 1**). First, promising ATAC-seq peaks were identified proximal to striatal cell type marker genes (**Figure 2A**) in multiple previously published and publicly available human and mouse bulk ATAC-seq datasets (**Figure 2B**). The majority of these enhancers were selected from the human Brain Open Chromatin Atlas (BOCA^35^) and mouse Cis-element ATlas (CATlas^36,37^) as sequences with relatively strong brain region or cell type selectivity in chromatin accessibility (**Figure 2C-D**). An additional handful of enhancer candidates were selected from mouse cortical scATAC-seq data^21^ or from a published human M1 dataset^20^. These enhancers were found to exhibit useful cell type specificity in the striatum region in addition to cortex based on prior brain wide expression analysis. Additionally, virtually none of the selected enhancers exhibited accessibility in human tissues outside of the brain (**Figure S1**).

**Figure 1.**
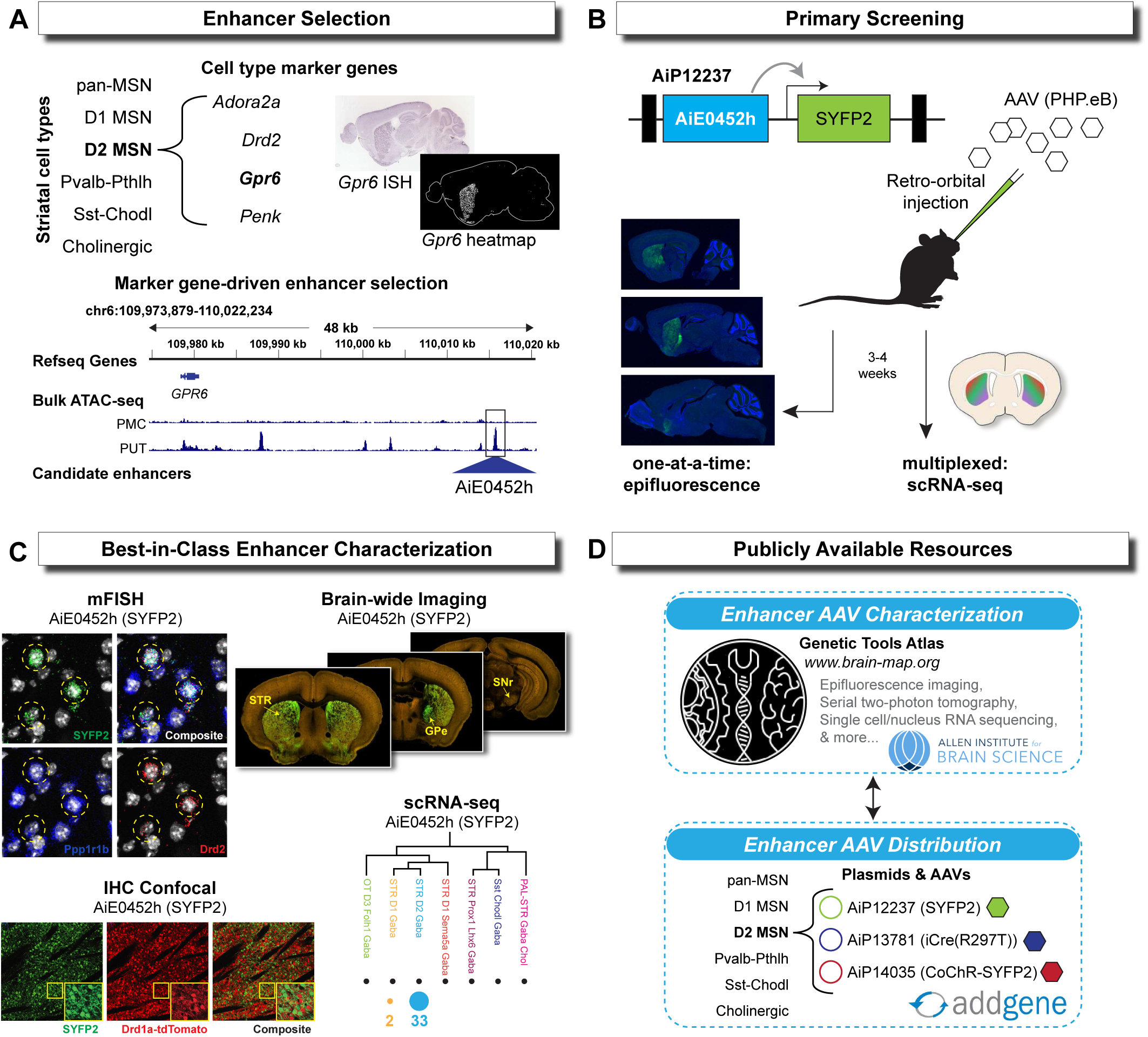
Pipeline for the discovery, validation, and distribution of striatal enhancer AAVs. A) Diagram of enhancer selection process. Isolated peaks are chosen based on proximity to marker genes from multiple different ATAC-seq datasets. B) Putative enhancer sequences were screened for cell type specific activity by retro-orbital injection using SYFP2 native fluorescence. Example shown is of plasmid AiP12237 that contains the D2 MSN enhancer AiE0452h driving expression of SYFP2. C) The most promising “Best-in-Class” enhancers were further validated for on-target activity by comparing mRNA and protein expression across multiple techniques. D) Striatal enhancer AAV characterization data is publicly available through the Allen Institute for Brain Science Genetic Tools Atlas (https://portal.brain-map.org/genetic-tools/genetic-tools-atlas). Enhancer AAV plasmid DNA and select virus aliquots are available from Addgene for distribution to the research community.

**Figure 2.**
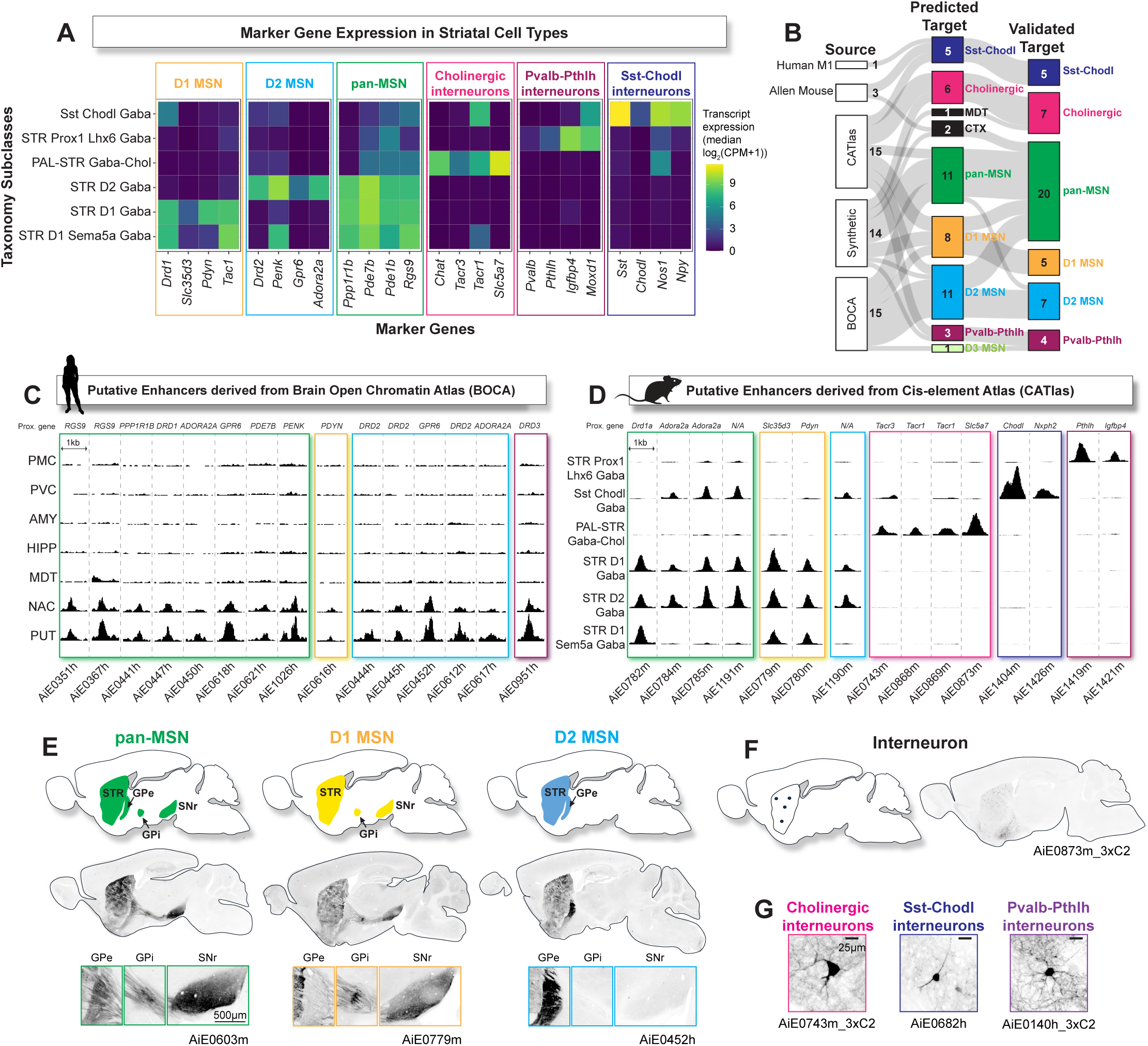
Putative enhancers can drive cell type specific expression in the striatum. A) Heatmap of median transcript detection log2(CPM+1) of striatal cell type marker genes from CCN20230722 mouse whole brain taxonomy cluster matrix. Colored boxes group marker genes by corresponding cell type common name. B) Sankey diagram with three nodes indicating the sources for all putative enhancer peaks for each cell target, their predicted expression specificities, and their validated expression. Values indicate number of enhancers at each node. C-D) Putative enhancer peaks selected for striatal cell types from BOCA (C) and CATlas (D) datasets. Tracks are grouped by brain region for BOCA (PMC: primary motor cortex, PVC: primary visual cortex, AMY: amygdala, HIPP: hippocampus, MDT: medio-dorsal thalamus, NAC: nucleus accumbens, PUT: putamen) and by CCN20230722 whole mouse brain taxonomy designated cell subclass for CATlas. The marker gene each peak was found proximal to is noted above each peak (N/A indicates peak was not found near any known marker gene). Enhancer peaks are grouped by target (gold: D1 MSN, turquoise: D2 MSN, green: pan-MSN, pink: cholinergic interneurons, dark blue: Sst-Chodl interneurons. Maroon: Pvalb-Pthlh interneurons. E) Top: diagram of expression patterns indicative of striatal MSN cell populations. Middle: example sagittal sect ions of native SYFP2 expression from enhancer AAVs for each indicated MSN population. Bottom: zoomed in views of axonal projection patterns in basal ganglia target regions from the same enhancer AAV as middle panel (GPi: globus pallidus internal segment, GPe: globus pallidus external segment, SNr: substantia nigra pars reticulata). F) Left: diagram of expression patterns indicative of striatal interneuron populations. Right: example sagittal section of native SYFP2 from enhancer AAV with interneuron expression selectivity. G) Example of morphological differences for distinguishing striatal interneuron populations. Note large size of cholinergic interneuron somata and thick dendrite caliber, elongated or bipolar Sst-Chodl interneurons with thin dendrite caliber, and compact dendrites of multipolar Pvalb-Pthlh interneurons.

Candidate enhancer sequences were PCR amplified from mouse or human genomic DNA and cloned into an AAV vector upstream of a minimal promoter sequence (either minimal beta globin or minimal Rho promoters) and the bright monomeric SYFP2 fluorescent reporter transgene^38^. Completed vectors were packaged into functional AAV particles using the mouse blood brain barrier penetrant capsid PHP.eB^39^ and injected retro-orbitally (RO) into adult wildtype C57Bl6/J mice. At 3-5 weeks post injection the mouse brains were collected for analysis of native SYFP2 reporter expression pattern in the brain.

As striatal medium spiny neurons (MSNs) make up ∼95% of all striatal neurons and are defined by their axon projection targets^40,41^, D1 MSN and D2 MSN selective labeling was easily identifiable in mid-sagittal brain sections as dense SYFP2^+^ striatal neuron labeling with axons projecting predominantly to the GPi/SNr or GPe, respectively (**Figure 2E**). Similarly, pan-MSN labeling was revealed as dense SYFP2^+^ striatal neuron labeling with axons projecting to both direct and indirect pathway target structures. Several of the MSN enhancers exhibited intra-striatal regional biases, with cell body expression mostly restricted to either dorsal or ventral striatum (**Figure S2**). In contrast, enhancers that labeled local interneuron populations exhibited SYFP2 fluorescence restricted to the striatum proper and had clearly distinguishable morphological characteristics (**Figure 2F-G**)^5,14,42^ ^44^. Striatal Pvalb-Pthlh interneurons have small round or rectangular somata with compact multipolar aspiny dendrites, cholinergic interneurons have unusually large cell bodies with thick aspiny dendrites, and Sst-Chodl interneurons have elongated small to medium size somata with wispy, thin dendrites.

Following initial evaluation by native SYFP2 fluorescence, we sought to improve enhancer-driven transgene expression by inserting multiple copies of the “core” enhancer region in tandem. This strategy has been previously shown to improve strength of expression while preserving specificity^21,45^. We designed and screened an additional 13 of these “synthetic” enhancers that resulted in stronger SYFP2 fluorescence compared to the original full-length enhancers (**Figure S3**).

Based on analysis of the striatum and brain wide expression patterns, we curated a set of 48 enhancer AAV vectors with promising activity spanning these major striatal neuron populations. This resulted in a collection of 20 pan-MSN enhancers, 5 D1 MSN enhancers, 7 D2 MSN enhancers, 5 Sst-Chodl interneuron enhancers, 7 cholinergic interneuron enhancers, and 4 Pvalb-Pthlh interneuron enhancers (**Figure 2B and S3; Table S1**). Most enhancers in this set showed high selectivity for the striatum, with minimal or no SYFP2^+^ cell labeling detected in other brain regions. In the case of Sst-Chodl or Pvalb subclass enhancers we often observed labeled neurons in both cortex and striatum regions.

### Comparison of strength and specificity of validated striatal MSN enhancers

In the case of enhancer selection based on bulk human ATAC-seq data from BOCA^35^, putative D1 MSN or D2 MSN enhancer candidates were inferred based on proximity to known direct and indirect pathway MSN marker genes (e.g., *Drd1* and *Drd2*)^12^. Additional criteria included differential chromatin accessibility in striatum neuronal samples versus all other brain region neuronal samples including multiple cortex regions, thalamus, hippocampus, and amygdala. Because the source epigenetic data used for selection did not have the resolution of subclasses we were targeting, we found that some enhancers inferred as putative D1 MSN or D2 MSN enhancers instead yielded pan-MSN reporter activity that was nonetheless fully consistent with the bulk epigenetic signatures.

To explore the relative strength of a set of validated pan-MSN enhancers, we designed a multiplex experiment to directly compare the enhancers in the mouse brain *in vivo* (**Figure S4A**). We chose to include twelve pan-MSN enhancers as well as one D1 MSN enhancer and one D2 MSN enhancer to confirm our ability to resolve subclass specific labeling. An 8-base pair (bp)-barcode was cloned between the WPRE3 and bGHpA of each parental enhancer AAV construct, packaged separately into PHP.eB and administered RO into an adult mouse as a single pooled injection. SYFP2^+^ whole cells were collected via FACS and used to generate 10x v3.1 single cell gene expression and AAV transcript (barcode) libraries. Enhancer specificities were determined by the presence of an AAV barcode following mapping of sequenced libraries to the mouse whole brain taxonomy (see **Methods** for details).

We examined properties that reflect enhancer cell type specificity, strength, and broadness of labeling across the striatum. We did not observe any major differences in cell type specificity, as almost all pan-MSN enhancer AAVs labeled equivalent proportions of D1 and D2 MSNs (STR D1 Gaba and STR D2 Gaba in the taxonomy). The exception was AiE0450h which exhibited a STR D2 Gaba bias, which is relatively consistent with its initial selection as an *ADORA2A*-proximal enhancer candidate (**Figure S4B**). The subclass specific enhancers AiE0779m_3xC2 and AiE0452h_3xC2 had greater than 90% specificity for their respective cell types (90.4% STR D1 Gaba and 92% STR D2 Gaba, respectively, FIG). We observed a dynamic range in number of cells expressing enhancer-driven transcripts, with the weakest enhancer-barcode detected in 250 cells (AiE0526h) to the strongest in 2,067 cells (AiE0784m) (**Figure S4B**), suggesting large differences in completeness of striatal MSN labeling in the dorsal striatum region between enhancers. We also counted the number of barcoded transcripts per cell as a measure of enhancer strength and used the Kruskal-Wallis rank sum test to assess differences (chi-squared = 1163.3, df = 11, p < 0.0001) (**Figure S4C**). Dunn’s post hoc analysis revealed significant differences between many of the enhancers. Importantly, both full length enhancers, AiE0441h and AiE0367h, had significantly fewer transcripts detected than the optimized versions, AiE0441h_3xC2 and AiE0367h_3xC2 (AiE0367h vs AiE0367h_3xC2: Z = −7.43, p <0.0001 and AiE0441h vs AiE0441h_3xC2: Z = −17.30, p <0.0001), further validating enhancer core concatenation improved enhancer performance. The two strongest enhancers, AiE0784m and AiE0441h_3xC2, exhibited both broad coverage as well as significantly higher transcript expression compared to all other enhancers in the dorsal striatum.

Native SYFP2 fluorescence in sagittal sections following individual retro-orbital (RO) injection of the barcoded enhancer AAV vectors (**Figure S4D-E**) provided additional independent corroboration of the *in vivo* multiplex results. The dynamic range in native SYFP2 fluorescence intensity closely matched both the multiplex counts and primary screening results using the parental vector (**Figure S2**). These results confirm AiE0441h_3xC2 and AiE0784m as our top two strongest pan-MSN enhancer AAVs.

### Multimodal molecular validation confirms striatal enhancer AAV cell type specificity

We next sought to quantitatively measure enhancer specificity in the dorsal striatum by employing a combination of immunohistochemistry (IHC), single cell RNA sequencing (scRNA-seq), and RNAscope (**Figure 3A**). Using IHC for protein detection, we analyzed native enhancer-driven SYFP2 fluorescence for 15 enhancers together with striatal cell type marker antibodies (anti-ChAT for cholinergic interneurons, anti-Pvalb for Pvalb-Pthlh interneurons, and anti-nNos for Sst-Chodl interneurons^15,34^).

**Figure 3.**
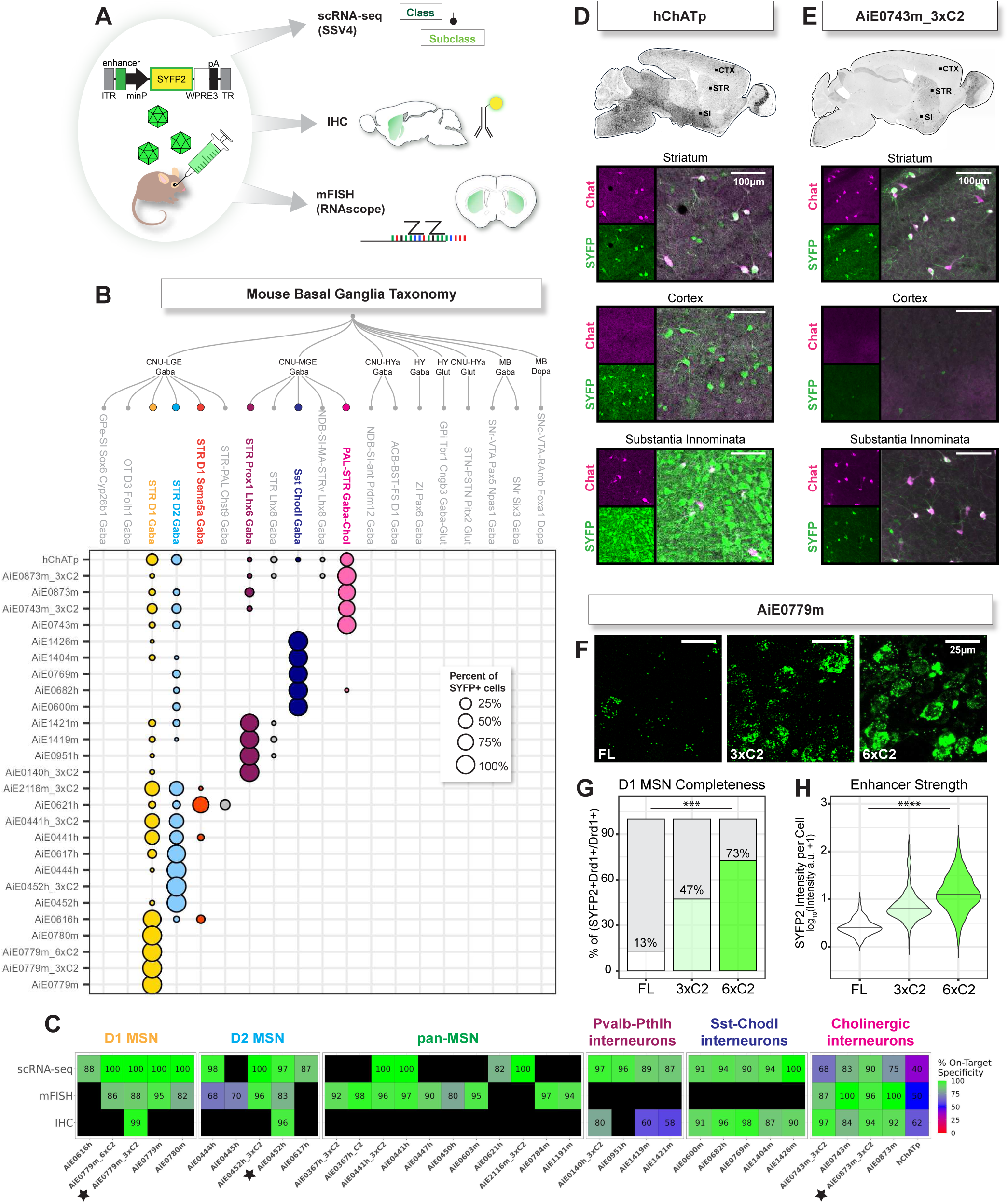
Striatal enhancer AAVs demonstrate significant improvement in cell type specificity over existing tools. A) Diagram of triple modality enhancer AAV validation process including scRNA - seq (Smart-seq V4, SSv4), immunohistochemistry (IHC), and mFISH (RNAscope). B) Enhancer specificities determined by SSv4 scRNA-seq (n=1-2 mice and on average 50 cells per enhancer). Sequenced transcriptomes were mapped against the 10X Whole Mouse Brain taxonomy CCN20230722 using hierarchical mapping algorithm in MapMyCells (Allen Brain Map). Cell types in dendrogram encompass all neuronal basal ganglia subclasses from the taxonomy, with the six cell types targeted in this manuscript highlighted in color. C) Quantification of on-target enhancer AAV specificities across all three validation methods. Black boxes indicate no data and stars indicate 3 of the top enhancers. D-E) Top: example sagittal sections of hChATp (D) or AiE0743m_3xC2 (E) driven native SYFP2 fluorescence. Bottom: Identification of striatal cholinergic interneurons by IHC using ChAT subclass marker antibody. Each panel is a representative ROI from three different brain regions. F) RNAscope using probes against SYFP2 for three versions of the AiE0779m D1 MSN enhancer (FL=full length enhancer, 3xC2=optimized enhancer with 3x core region, and 6xC2=optimized enhancer with 6x core region). G) Quantification of completeness of labeling D1 Fisher’s exact test revealed differences in proportions of SYFP2-positive cells between AiE0779m enhancer versions (*p* < 0.0001). Post-hoc pairwise comparisons with Bonferroni correction showed significant differences between all enhancers (\*\*\**p* <0.001). H) SYFP2 signal intensity per cell from RNAscope images. The Kruskal-Wallis test revealed a significant difference in SYFP2 intensity across enhancers X^2^ = 424.39, df = 2, *p* < 0.0001). Post-hoc Dunn’s test with Bonferroni correction showed that all pairwise comparisons were significant: 3xC2 vs. 6xC2 (*p* < 0.0001), 3xC2 vs. FL (*p* < 0.0001), and 6xC2 vs. FL (*p* < 0.0001). n=1-2 mice and 1 ROI per enhancer version for RNAscope quantification.

Enhancers specific for striatal D1 and D2 MSN subclasses were analyzed using SYFP2 fluorescence following RO virus injection into Drd1a-tdTomato transgenic mice and confocal imaging. To resolve the identity of enhancer AAV labeled striatal cell types, single cell RNA sequencing was performed on 27 enhancers using Smart-seq v4 (SSv4) on isolated SYFP2^+^ whole cells dissociated from the dorsal striatum region. We collected and sequenced on average 50 cells per experiment (one enhancer AAV vector RO injected into one mouse; average cells +/-SD = 50 +/- 15.95) for a total of 1,342 cells. Cell subclasses were defined by mapping the cells to the Allen Institute’s mouse whole brain taxonomy using the MapMyCells hierarchical mapping (https://portal.brain-map.org/^30^). (**Figure 3B**). Lastly, we performed multiplex fluorescent in situ hybridization (mFISH) using RNAscope for 22 enhancer AAVs using probes against SYFP2 and well-known striatal cell type marker genes *Ppp1r1b* (pan-MSNs), *Drd1* (D1 MSNs), *Drd2* (D2 MSNs), and *Chat* (cholinergic interneurons) (**Figure S5**).

We compared the specificity of each enhancer AAV calculated from all three modalities (**Figure 3C**). In general, all enhancers exhibited high specificity for their designated cell subclass (overall specificity across techniques +/- SD = 88.5% +/- 13%) and all techniques were mostly in agreement (IHC average specificity +/- SD =85.57% +/- 13.86%, SSv4 average specificity +/- SD=90.58% +/- 13.21%, RNAscope average specificity +/- SD =88.13% +/- 12.81%). For our exemplary D1 and D2 MSN enhancers we obtained specificity values across modalities that were in close agreement in support of very high specificity at the subclass level. For D1 MSN enhancer AiE0779m_3xC2 the specificity was calculated as 99% by IHC (SYFP2^+^tdTomato^+^/SYFP2^+^) and 100% by scRNA-seq and taxonomy mapping. For D2 MSN enhancer AiE0452h the specificity was 96% by IHC (SYFP2^+^tdTomato^-^/SYFP2^+^) and 97% by scRNA-seq and mapping. In both cases the specificity was slightly lower by mFISH/RNAscope (88% and 83% subclass specificity, respectively), although still in close agreement. Slightly lower values could be due to high noise in mFISH cell segmentation and image analysis or the preferential collection of bright cells by FACS in scRNA-seq.

A relatively larger disparity in specificity measurements between techniques was found for enhancers labeling the Pvalb-Pthlh interneuron subclass, where specificity values determined by SSv4 was significantly higher than by IHC (Welch Two Sample t-test p=0.047; SSv4 average +/- SD =92.33% +/- 4.41% vs. IHC average +/- SD =65.83% +/- 9.7%). This aligns with recent evidence indicating that *Pvalb* expression is found in many but not all interneurons belonging to this striatal neuron subclass^28^. Accordingly, we report that the SSv4 approach is better suited for determining the specificity of striatal Pvalb-Pthlh subclass enhancers, since this technique utilizes the collective transcriptome for mapping as opposed to relying on a single marker gene. With this in mind, we validated four different striatal Pvalb-Pthlh interneuron enhancers with specificity ranging from 87-97% at the subclass level by SSv4. We also validated five novel striatal Sst-Chodl interneuron enhancers with specificity ranging from 87-100% at the subclass level and with close agreement between IHC and SSv4 (**Figure 3C**), thus mFISH analysis was not deemed necessary.

We cloned a previously identified 894 bp human *Chat* promoter fragment (hChATp) into our AAV vector design for comparison to our enhancers for labeling striatal cholinergic interneurons. This human *Chat* promoter fragment was previously used in canine adenoviral (CAV) and AAV vectors to achieve enriched labeling of cholinergic neurons following stereotaxic injection into the mouse, rat and macaque striatum with specificity ranging from 64-86% for ChAT^+^ neurons^46,47^. Here, we tested the hChATp for brain wide specificity of labeling in mouse following RO delivery in our platform. Four of our novel cholinergic enhancers achieved higher specificity of labeling in the dorsal striatum by the three quantification methods (**Figure 3C**) and with lower off-target expression in cortex, substantia innominata, and hindbrain regions, as illustrated by ChAT co-staining images for AiE0743m_3xC2 relative to hChATp (**Figure 3D-E**). Our best-in-class striatal cholinergic enhancer AiE0873m_3xC2 achieved a consensus of >90% specificity at the subclass level by all three quantification methods (IHC, SSv4, and mFISH), as compared to a highest value of 62% specificity for hChATp by IHC.

In addition to quantifying specificity of labeling we also measured signal intensity in RNAscope experiments to compare SYFP2 reporter expression levels between the full length AiE0779m enhancer with the optimized versions, AiE0779m_3xC2 and AiE0779m_6xC2, extending our previously established enhancer core concatenation approach^21,22^. We found the number of concatenated enhancer cores was positively correlated with both enhancer strength and completeness of labeling (**Figure 3F-H**). Although previous work has demonstrated that concatenation of multiple enhancers proximal to a given gene of interest does not generally appear to be an efficient method for identifying highly specific enhancers^48^, future work employing DNA sequence-based models of enhancer activity could generate novel synthetic enhancers that could greatly refine cell type-specific targeting efforts.

We also used scRNA-seq datasets to assess expression levels of marker genes following enhancer AAV viral transduction across all five striatal cell types. Because enhancer AAVs use transcription factors (TF) to activate gene expression, they may compete with endogenous enhancers for the same TFs, “enhancer squelching^49^”. To test this possibility, we examined the expression of marker genes in cell populations targeted by AAVs containing enhancers proximal to that marker gene versus populations targeted by other enhancers. Overall, we found marker gene expression in proximal vs non-proximal enhancer AAVs to be very similar (R^2^=0.9) suggesting most enhancers do not exhibit TF squelching (**Figure S6**). Only the gene *Adora2a* exhibited lower average expression in cells labeled by the D2 MSN proximal enhancer AiE0617h. However, it remains unclear whether the observed reduction in *Adora2a* expression in cells labeled by AiE0617h is caused by TF squelching or reflects natural variation in *Adora2a* expression across D2 MSN cell subtypes. Regardless, this reduction was only observed for a single enhancer and thus does not appear to be a universal effect.

### Comparison of striatal enhancer performance by different routes of virus administration

It was previously reported that stereotaxic injection of neocortical cell type enhancer AAVs led to lower cell type specificity values as compared to RO injection of the same vectors^21^. Given the relative dearth of evidence along these lines, it is important to evaluate enhancer specificity for different routes of AAV administration and to document the optimal dose ranges associated with each route (**Figure 4A**). We tested a subset of exemplary striatal cell type enhancers driving SYFP2 reporter transgene by three different routes of administration (RO, intracerebroventricular (ICV), and stereotaxic injection) and compared the specificity of labeling (**Figure 4B-E**). We qualitatively evaluated specificity by analysis of direct versus indirect pathway axon projection targets as well as performed scRNA-seq on SYFP+ cells from the dorsal striatum (**Figure 4F-G**). D1 MSN enhancer AiE0779m drove on-target SYFP2 expression by RO and ICV routes but exhibited an apparent loss of specificity by stereotaxic injection based on dense axon terminal labeling observed in GPe in addition to GPi and SNr (**Figure 4B**), despite multiple rounds of iterative adjustment to AAV dosage (data not shown). However, scRNA-seq analysis demonstrated AiE0779m still maintained 100% specificity for D1 MSNs across routes (**Figure 4F-G**). Therefore, the observed signal in GPe may instead be axon collaterals of D1 MSNs terminating in GPe in addition to GPi and SNr^50,51^. In contrast, clear on-target labeling of direct pathway D1 MSNs was observed for AiE0780m by all routes of AAV delivery (**Figure 4C**) and corroborated by scRNA-seq (**Figure 4F-G**). Similarly, exemplary D2 MSN enhancer AiE0452m yielded strong on target SYFP2 expression in direct pathway D2 MSNs with all routes of administration (**Figure 4D**).

**Figure 4.**
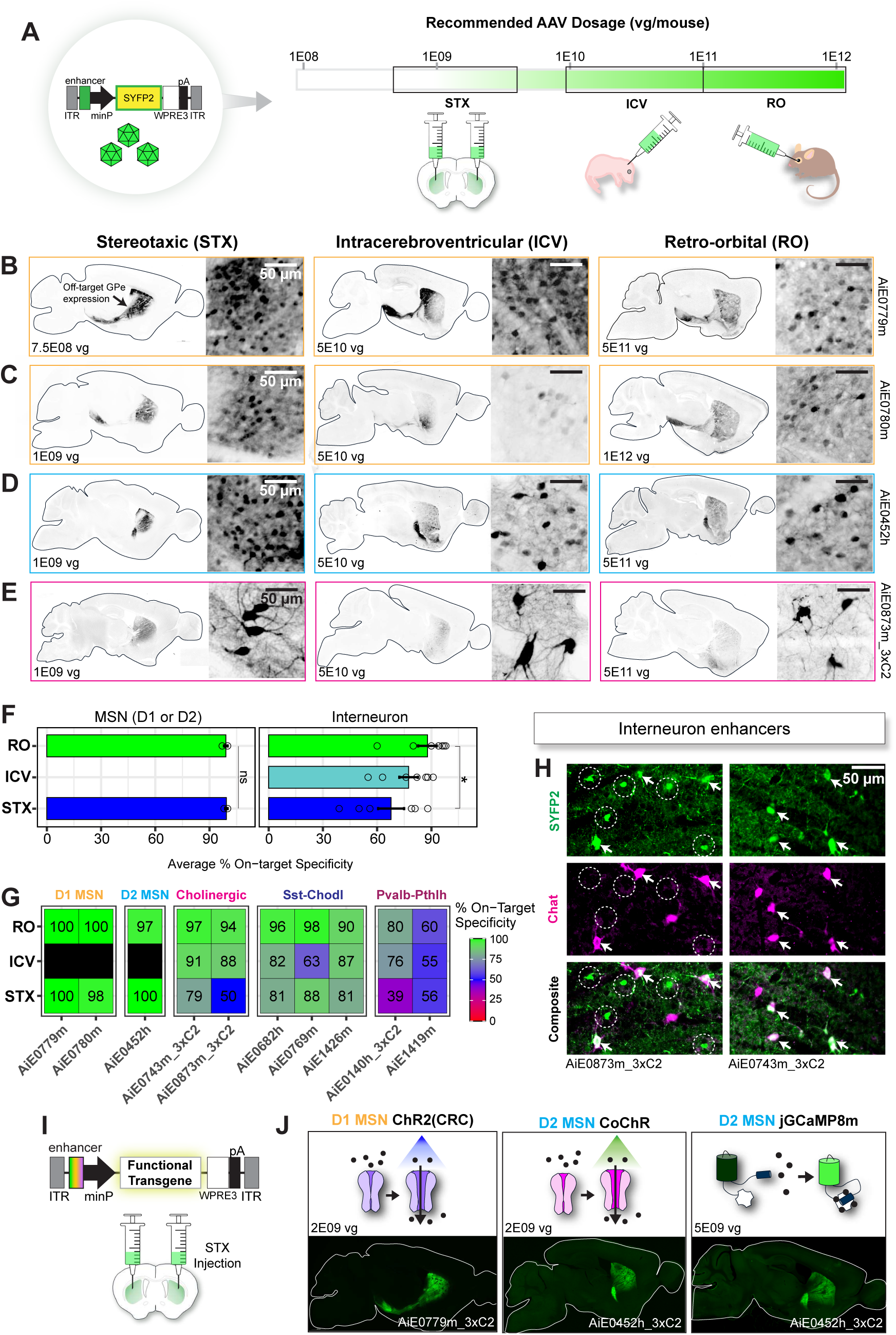
Specificity of labeling for striatal enhancer AAVs across different delivery routes and cargos. A) Diagram of enhancer AAV vector build and range of recommended AAV dosages across stereotaxic (STX), intracerebroventricular (ICV), and retro-orbital (RO) routes of administration in mice. Note ICV delivery is in P1-P2 pups while STX and RO are in adult mice. B-E) Representative sagittal sections of native SYFP2 fluorescence across all three delivery routes for four dif ferent enhancers targeting B-C) D1 MSNs, D) D2 MSNs, and E) cholinergic interneurons. F) Summary of D1 and D2 MSN enhancer and interneuron enhancer specificities by route of administration. Each point represents a single enhancer. A Kruskal-Wallis test revealed no significant difference in specificity across routes for the MSN enhancers (*p* = 0.7963), while a significant difference was observed for interneuron enhancers (*p* = 0.04383). Post-hoc Dunn’s test with Bonferroni correction indicated that the RO route dif fered significantly from the STX route (*p* < 0.05), but no signif icant difference was found between ICV and other routes. G) Heatmap of on-target specificities for striatal enhancers for the three routes of administration determined by scRNA-seq and mapping (D1 and D2 MSN enhancers) or IHC with cell type marker antibodies (interneuron enhancers). H) Images of IHC comparing two cholinergic enhancers AiE0873m_3xC2 and AiE0743m_3xC2 by STX injection. White arrows show colocalization of SYFP2 and Chat. White circles in the AiE0873m_3xC2 images demonstrate off-target SYFP2 expression in MSNs. I) Diagram of enhancer AAV vector build for delivery of functional transgenes by STX injection into dorsal striatum. J) Representative sagittal images of native fluorescence for enhancer AAVs driving functional transgenes. Left: D1 MSN enhancer AiE0779m_3xC2 driving ChR2(CRC)-EYFP (AiP13278). Middle: D2 MSN enhancer AiE0452h_3xC2 driving CoChR-EGFP (AiP14035). Right: D2 MSN enhancer AiE0452h_3xC2 driving jGCaMP8m (AiP14134).

For striatal interneuron subclass enhancers (cholinergic: AiE0873m_3xC2 and AiE0743m_3xC2; Pvalb-Pthlh: AiE0140h_3xC2; Sst-Chodl: AiE0769m and AiE0682h) we performed immunostaining with cell type marker antibodies (ChAT, Pvalb, and nNos, respectively) and quantified specificity of labeling in the target subclass in the dorsal striatum region by each route of virus administration (**Figure 4F-G**). In all cases the specificity of labeling was highest with RO injection (range of 80%-98% specificity) and lowest with stereotaxic injection (range of 39%-88% specificity). The ICV injection route produced intermediate specificity values (range 63%-91%). The lower specificity observed with stereotaxic injection versus RO injection for this set of striatal interneuron enhancer AAV vectors may indicate that further optimization of the injection dose is necessary, although specificity was likely at or near optimal for Sst-Chodl enhancer AiE0769m at 88% vs. 98% and cholinergic enhancer AiE0743m_3xC2 at 79% vs. 97% for stereotaxic vs. RO, respectively. For the second cholinergic enhancer AiE0873m_3xC2, the specificity of labeling by RO (94%) and ICV (88%) routes was exceptionally high but fell precipitously with stereotaxic injection route (50%). Examination of the dorsal striatum high magnification images revealed many smaller putative MSNs detected by antibody amplification of the SYFP2 signal, in addition to the large ChAT^+^ cholinergic neurons (**Figure 4H**).

### Versatility of striatal enhancers for expression of diverse transgene cargos

Similar to route of administration, the packaged transgene cargo could also impact vector expression level or specificity^52^. To directly address this possibility, we swapped out SYFP2 for several commonly used functional cargos including CoChR-EGFP and the calcium indicator jGCaMP8m for direct expression under the control of our best-in-class D1 and D2 MSN enhancers (**Figure 4I-J**). The packaged AAVs were administered into the dorsal striatum by stereotaxic injection (**Figure 4I**), a route frequently used for neural circuit dissection and qualitatively checked for specificity and expression by native fluorescence and evaluation of axon projection targets of the direct versus indirect pathway. Viral dose was titrated to determine the optimal dose for balancing intended cell type specificity with robust transgene expression level. In contrast to the optimal stereotaxic injection dose of 1E+9vg for D2 MSN expression of SYFP2 under the AiE0452h enhancer (**Figure 4D**), we found that the optimal dose for expression of CoChR-EGFP was two times higher at 2E+9vg and five times higher at 5E+9vg for jGCaMP8s, indicating that these transgenes are relatively more difficult to express in striatal MSNs than SYFP2. Nonetheless, the specificity of labeling was well-maintained, as judged by selective axon terminal labeling in the GPe with no axon terminal labeling seen in GPi or SNr regions in both cases (**Figure 4J**).

Another use case for our striatal enhancer AAVs is cell type-specific conditional gene deletion using Cre/loxP technology. Typically, this is achieved by crossing transgenic mouse lines with loxP flanked (i.e., ‘floxed’) exons or gene loci to transgenic Cre driver lines, where Cre recombinase is expressed under the control of endogenous cell type specific regulatory elements^53^. Similarly, Cre-dependent expression can be achieved by crossing cell type-specific Cre driver lines with Cre-dependent reporter lines (**Figure 5A**), such as Ai14 Cre-dependent tdTomato reporter line^54^. While there are several options for transgenic mouse lines driving Cre recombinase in striatal D1 and D2 MSNs^55,56^, many have extensive extra-striatal labeling (either consistent with endogenous gene expression patterns or ectopic in nature^11^) making them unsuitable for striatum-specific perturbations in disease modeling.

**Figure 5.**
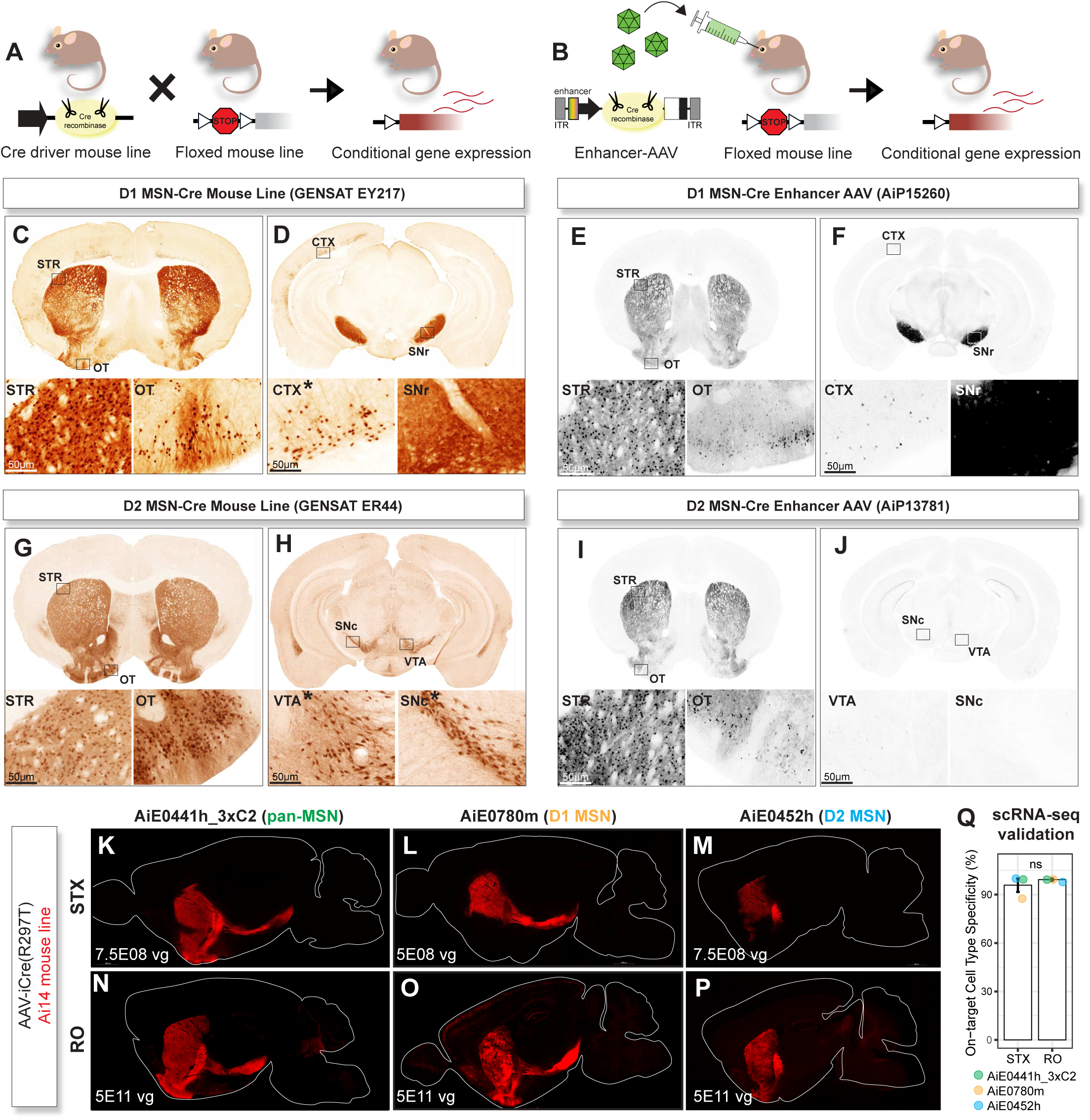
Comparison of D1 MSN and D2 MSN enhancer AAVs to GENSAT Cre driver lines. A) Diagram of breeding cross for generating conditional reporter transgenic mice by Cre/loxP recombination. B) Diagram of strategy for generating conditional reporter transgene expression using enhancer AAVs. C-D) ISH from www.gensat.org of GENSAT EY217 Drd1a-Cre driver line crossed to Rosa26-EGFP reporter line with C) cell body expression in striatum (STR) and islands of calleja in olfactory tubercle (OT) and D) projections to substantia nigra pars reticulata (SNr) with extra-striatal expression observed in cortex. E-F) STPT images of D1 MSN enhancer AiE0779m driving iCre recombinase point mutant R297T (AiP15260) injected into Ai14 reporter mouse line with E) cell body expression in STR but not islands of calleja of the olfactory tubercle (OT) and F) projec tions to SNr. G-H) ISH from www.gensat.org of GENSAT ER44 Drd2-Cre driver line crossed to Rosa26-EGFP reporter line with G) cell body expression in STR but not islands of calleja in OT and H) extra-striatal dopamine neuron expression in ventral tegmental area (VTA) and SNr. I-J) STPT images of D2 MSN enhancer AiE0452h driving iCre recombinase point mutant R297T (AiP13781) injected into Ai14 reporter mouse line with I) cell body expression in STR but not islands of calleja in OT and J) no off-target midbrain dopamine expression. K-P) Example sagittal sections of enhancer AAVs driving iCre(R297T) (AiP14825, AiP15578, and AiP13781) in Ai14 reporter mouse line either through STX (K-M) or RO (N-P) injection. Images are of native tdTomato fluorescence. Q) Comparison of on-target cell type specificity using scRNA-seq between RO and STX injection methods. Bars represent mean ± standard error (SE). Individual data points indicate values for each enhancer. A Wilcoxon rank sum test was performed to assess statistical differences between groups (W = 5, *p* = 1), indicating no significant difference between injection methods.

Two of the most specific and widely used mouse Cre driver lines are the EY217 (D1 MSN) and ER44 (D2 MSN) GENSAT BAC transgenic mice^11^. We compared EY217 and ER44 mouse line brain expression data from the GENSAT website (www.gensat.org) with two of our best-in-class MSN enhancers, AiE0779m_3xC2 (D1 MSN) and AiE0452h (D2 MSN), driving attenuated iCre(R297T), a point mutant iCre variant that was successful at improving the fidelity of cell type specific enhancer recombination mitigating extra-striatal labeling^25,52^. The enhancer AAVs were delivered by RO injection into Ai14 Cre-dependent tdTomato reporter mice (**Figure 5B**). We observed several improvements in the enhancer AAV strategy compared to the GENSAT Cre lines. First, the D1 Cre BAC Tg mouse line EY217 exhibits expression in granule cells of the islands of Calleja in the olfactory tubercle (OT) and in layer 6 of the neocortex (**Figure 5C-D**), whereas far less labeling was seen in these regions and cell types with D1 MSN enhancer Cre driven recombination (**Figure 5E-F**). Second, in the case of D2 Cre BAC Tg mouse line ER44, there is expression of Cre recombinase in both striatal D2 MSNs and in midbrain dopamine neurons known to express the D2 receptor and that send dense axon projections to the striatum (**Figure 5G-H**). In contrast, the D2 MSN enhancer driven Cre recombination was more striatum-restricted in D2 MSNs with no expression observed in midbrain dopamine neurons (**Figure 5I-J**). Thus, our best-in-class D1 and D2 MSN Cre AAV vectors compare favorably to widely used D1 and D2 Cre BAC transgenic mouse lines.

Cre recombination exhibits high sensitivity and can be challenging to titrate under certain experimental contexts such as stereotaxic injection where the effective multiplicity of AAV infection is much higher compared to RO injection. We selected pan-MSN enhancer AiE0441h_3xC2, D1 MSN enhancer AiE0780m, and D2 enhancer AiE0452h to pair with iCre(R297T) in AAV vectors for testing cell type specific Cre recombination by stereotaxic injection into the dorsal striatum versus RO injection in adult Ai14 mice (**Figure 5K-P**). We titrated the optimal dose for stereotaxic injection experiments and achieved highly specific Cre recombination in MSNs, D1 MSNs, and D2 MSNs. The specificity of labeling was quantitatively comparable (Wilcoxon rank sum test, p=1) between stereotaxic injection and RO injection experiments at the respective optimal doses by scRNA-seq measurements (**Figure 5Q**). Using this same strategy, we generated iCre(R297T) and FlpO vectors for successful and highly specific cell type recombination across all major MSN and interneuron cell types in the striatum (**Figure S7**). Additional scRNA-seq analysis revealed that specificity was well-maintained and not significantly different comparing SYFP2 vs. iCre(R297T) vectors across the collective subclasses, with on target specificity >90% for all but one Pvalb-Pthlh iCre(R297T) vector at 80.5% (**Figure S7B**).

### Enhancer driven ChR2 expression is sufficient for robust optogenetic perturbation in acute mouse brain slices and in vivo

An important question is whether cell type enhancers can drive meaningful levels of transgenes such as ChR2 to robustly control neuronal function and animal behavior with light. We first evaluated optogenetic stimulation using the acute mouse brain slice preparation following STX injection of AAV vectors into the dorsal striatum region in adult mice for targeted expression of ChR2 in D1 MSNs. We chose the ChR2(CRC) variant which includes mutations L132**C**/H134**R**/T159**C** and was shown to confer strong blue light evoked currents as described previously^57^. We compared injection of AiP13278, which utilizes D1 MSN enhancer AiE0779m_3xC2 for ChR2(CRC)-EYFP expression in C57 mice, versus injection of AAV-hEF1a-DIO-ChR2(CRC)-EYFP into D1-Cre transgenic mouse line. We additionally included testing of vector AiP15578 for D1 enhancer driven iCre(R297T) injected into dorsal striatum of the well-characterized Ai32 Cre-dependent ChR2(H134R)-EYFP mouse line^58^. we measured photocurrent generated by 1s blue light stimulation of recorded ChR2-expressing neurons in the dorsal striatum region (**Figure S8A)**. We found no significant difference in the blue light evoked peak photocurrent amplitudes measured form the three groups (**Figure S8B**, one way ANOVA, p=0.9) and no significant difference for any pairwise comparison between groups (Tukey’s multiple comparisons test, p>0.9 for all comparisons) (**Figure S8C)**. The large photocurrent amplitudes observed were sufficient to robustly and repeatedly drive action potential firing of D1 MSNs from rest for all three expression strategies (**Figure S8B)**.

We next sought to establish whether enhancer driven transgene expression is sufficient to perturb striatal neuron function *in vivo*. We delivered vector AiP13278 by ICV injection in P2 mouse pups for expression of ChR2(CRC)-EYFP in D1 MSNs (**Figure 6A**). As a control for virus injection and fluorescent protein expression alone we injected additional mice with vector AiP13044 for expression of SYFP2 in D1-MSNs using the same enhancer AiE0779_3xC2. Early postnatal virus injection by ICV route was most suitable to allow sufficient transgene expression time prior to optic fiber implantation surgery in young adult mice. Optic fibers for blue light delivery were implanted over the dorsal striatum (**Figure 6B**), and animals were used for behavioral analysis at 3-6 months old. *In vivo* optogenetic stimulation robustly induced locomotion and overt contralateral rotations in D1-ChR2 mice (n=8 mice), whereas no effect was observed with blue light stimulus in D1-SYFP2 animals (n=9 mice; **Figure 6C-I**). The blue light-induced contralateral rotations were robust across repeated trials for each session and across the entire cohort of D1-ChR2 mice. Thus, enhancer driven ChR2 in D1 MSNs of the dorsal striatum was sufficient for optogenetic control of mouse motor behavior.

**Figure 6.**
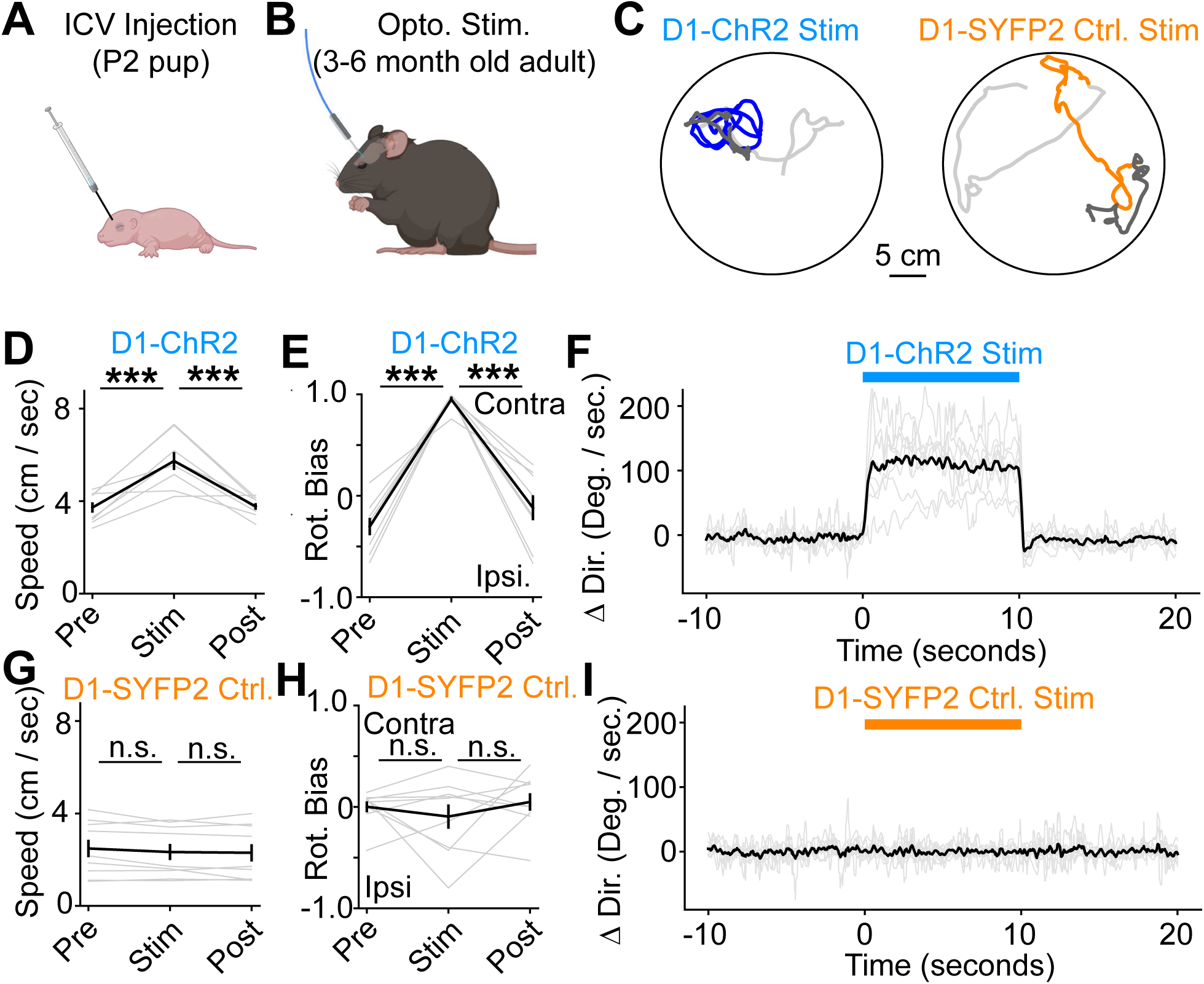
Optogenetic stimulation with viral vector targeting D1 MSNs is sufficient to induce locomotion and contralateral rotations. A-B) Experimental design with ICV injection of AAV vectors (AiP13278 for D1 ChR2 and AiP13044 for matched D1-SYFP2 control, 3.0E+10 vg each) into postnatal day 2 (P2) mouse pups. Optogenetic stimulation and behavioral tracking is performed at least 1 week after stereotaxic implantation of optical f ibers in adult mice. C) Exemplar tracks of mouse locomotion before (light gray; 10 sec), during (blue, orange; 10 sec), and after (gray; 10 sec) optogenetic stimulation in a 30 cm circular arena. D-E) Unilateral light delivery through the optical f iber (450 nm, 0.3 mW, 10 sec) increased the locomotion speed in D1-ChR2 virus injected mice (***p<0.001 by paired t-test; n=8 mice) but not D1-SYFP2 controls (ns, not significant; n=9 mice). F-G) Unilateral light delivery increased rotation bias towards contralateral rotations in D1-ChR2 injected mice (***p< 0.001 by paired t-test; n=8 mice) but not D1-SYFP2 control mice (ns, not significant; n=9 mice). H-I) Optogenetic stimulation induced steady contralateral rotations within ∼200 ms of the onset of light stimulation in D1-ChR2 virus injected mice but not D1-SYFP2 only controls. Rotations persisted throughout the stimulation.

### Multiplex viral labeling enables targeted recordings from three distinct striatal interneuron subclasses in wild-type mice

Simultaneous viral genetic labeling and imaging of discrete striatal neuron populations using contrasting fluorophores would be highly advantageous for assaying cellular features in genetic disease models, gene deletion studies, or other paradigms where complex transgenic animal crosses are time and cost prohibitive. To demonstrate such feasibility, we performed multiplex injections of striatal cell type enhancer AAVs driving contrasting fluorophores in adult mice (**Figure 7A**). We first combined RO injection of best-in-class D1 MSN, D2 MSN, and cholinergic enhancer AAV vectors for expression of SYFP2, monomeric teal fluorescent protein-1 (mTFP1), and tdTomato respectively. We observed mutually exclusive labeling of the direct versus indirect striatal projection pathways of the basal ganglia circuit together with the less abundant large cholinergic interneurons (**Figure 7B**). We did not observe enhancer crosstalk for this combination of striatal cell type enhancers. We also tested RO injection with enhancer AAVs for labeling pan-MSNs (AiE0447h driving SYFP2) and oligodendrocytes (AiE0410m driving mTFP1) and observed mutually exclusive labeling (**Figure S9**).

**Figure 7.**
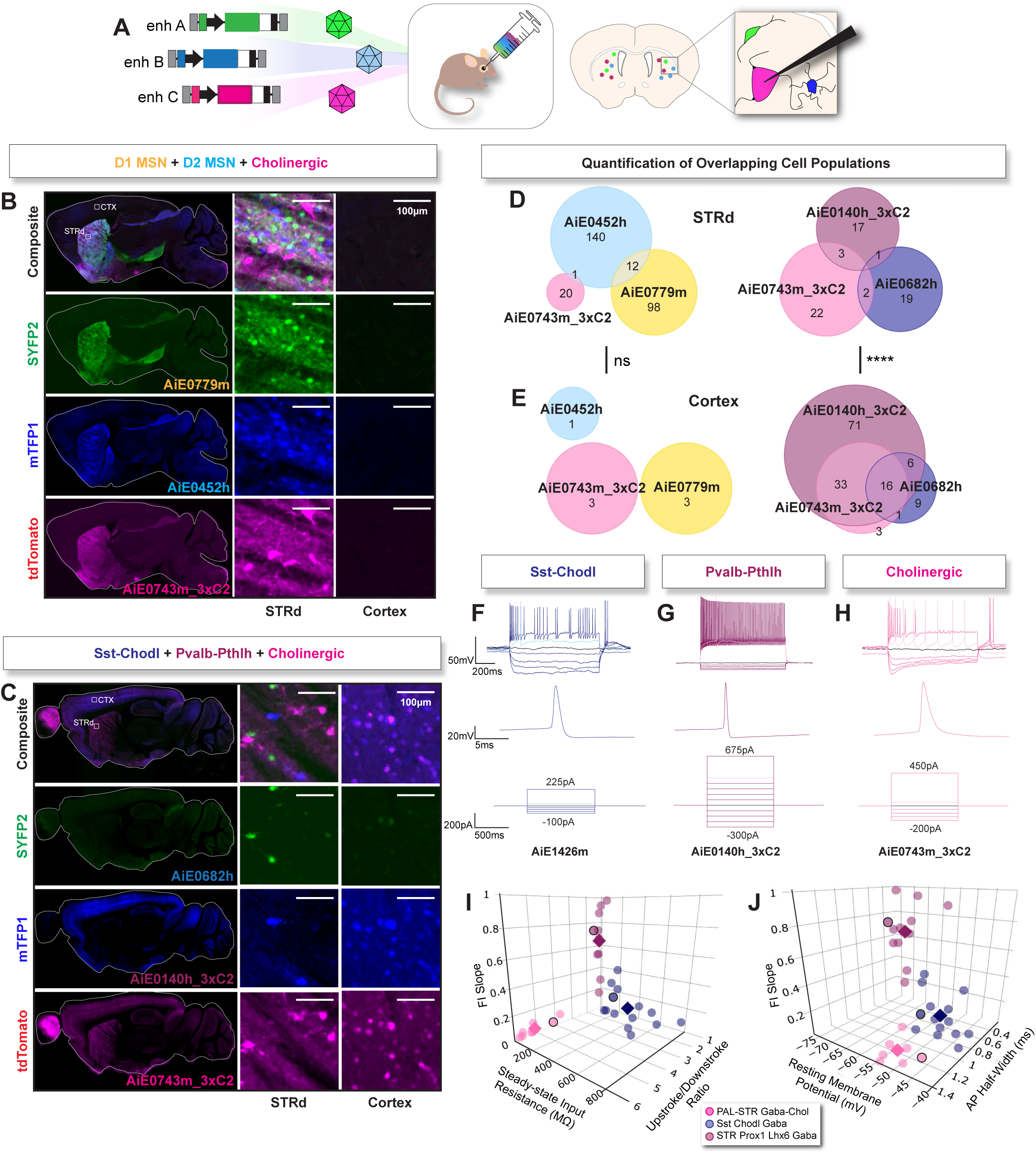
Concurrent striatal cell type identification and targeting by multiplexed enhancer AAV delivery in C57 mice. A) Enhancer AAVs were individually packaged, pooled, and injected RO into adult C57Bl6/J mice. B) Example sagittal sections of triple labeling using D1 MSN enhancer AiE0779m driving SYFP2 (AiP12609), D2 MSN enhancer AiE0452h driving mTFP1 (AiP12700), and cholinergic interneuron enhancer (AiE0743m_3xC2) driving tdTomato (AiP13738). Zoomed in views of dorsal striatum (STRd, middle) and cortex (right). C) Example sagittal sections of triple labeling using striatal Sst-Chodl enhancer (AiE0682h) driving SYFP2 (AiP12689), Pvalb-Pthlh enhancer (AiE0140h_3xC2) driving mTFP1 (AiP13808), and cholinergic enhancer (AiE0743m_3xC2) driving tdTomato (AiP13738). Zoomed in views of dorsal striatum (STRd, middle) and cortex (right). Note off-target tdTomato expression in cortex from AiE0743m_3xC2. D-E) Quantification of both multiplex injection combinations in dorsal striatum (D) and cortex (E). Numbers in Venn diagrams represent cell counts for each enhancer driven o evaluate enhancer expression overlap in cortex vs STRd (n=1 mouse and ROI per region). For D1 MSN+D2 MSN+Cholinergic enhancers, no significant dif ference was found (two-sided p = 0.8882) while for Sst-Chodl+Pvalb-Pthlh+Cholinergic enhancers, a significant difference in enhancer expression overlap between cortex vs STRd was observed (two-sided p <0.0001). F-H) Single cell patch-clamp recordings from coronal brain slices derived from a mouse with multiplex injection of Sst-Chodl enhancer AiE1426m-SYFP2 (AiP15050), Pvalb-Pthlh enhancer AiE0140h_3xC2-mTFP1 (AiP13808), and Cholinergic enhancer AiE0743m_3xC2-tdTomato (AiP13738). F-H): Representative voltage responses to hyperpolarizing and depolarizing current steps (top) with example single action potential traces highlighti ng distinctive AP halfwidths (middle). Current steps used to generate recordings are shown (bottom). I-J) 3D plots comparing electrophysiological features that distinguish the three striatal interneuron types. Individual neurons are shown as shaded circles, whereas darker diamonds represent the mean values for each interneuron type. Circles with black outline correspond to the exemplary neuron traces shown in panels F-H. PERMANOVA statistical testing using all f ive electrophysiological features found signif icant differences across the three cell type populations (F= 51.071, p= 0.001). A pairwise PERMANOVA post-hoc analysis indicated significant differences between all cell type pairwise comparisons (p<0.01). I) Comparison of the slope of the FI curve (f iring f requency vs. current injection), the steady state input resistance, and action potential upstroke to downstroke ratio. J) Comparison of the slope of the FI curve with the resting membrane potential and the action potential half-width. Sample sizes: Pvalb-Pthlh, n=13; Sst-Chodl, n=16; Cholinergic, n=8. A total of 7 mice were used for triple RO injection patch clamp recording experiments. Example voltage traces in panel F, G and H panels (top) were recorded from the same mouse.

Next, we performed a different triple RO injection combination to mark each of three major interneuron subclasses: Pvalb-Pthlh in mTFP1, Sst-Chodl in SYFP2, and cholinergic in tdTomato. Using the combination of AiE0140h_3xC2 (Pvalb-Pthlh), AiE0682h (Sst-Chodl), and AiE0743m_3xC2 (cholinergic) enhancers we uncovered surprising evidence of enhancer crosstalk in the neocortex.

Specifically, the AiE0743m_3xC2 cholinergic enhancer unexpectedly drove tdTomato expression in the same population of neurons marked by AiE0140h_3xC2 in the mTFP1 channel (**Figure 7C**). Notably, this pattern of widespread cortical tdTomato expression was not observed with AiE0743m_3xC2 enhancer AAV RO injection alone (**Figure S2**). Interestingly, the fluorescent protein expression patterns in the dorsal striatum region after the triple RO injection exhibited near mutual exclusivity, whereas the cortical region showed evidence of significant enhancer crosstalk (**Figure 7D-E**). To further explore this triple labeling paradigm for striatal interneuron types, we tested different combinations of novel enhancers for comparison to the preceding results. Using the combination of AiE0140h_3xC2 (Pvalb-Pthlh), AiE0743m_3xC2 (cholinergic), and alternate enhancer AiE1426m (Sst-Chodl), we no longer observed enhancer crosstalk in the brain. We instead observed exemplary mutually exclusive three-color labeling in the three distinct populations of striatal interneurons (**Figure S9F-J**).

We next sought to show functional analysis of cellular properties by targeted patch clamp recordings (**Figure 7F-J**). We targeted the three distinct striatal interneuron types for recordings based on their respective cytosolic fluorescent protein labels and their hallmark somatodendritic morphologies (**Figure 2G**). The tdTomato^+^ cholinergic interneurons and SYFP2^+^ Sst-Chodl interneurons both exhibited spontaneous action potential firing (data not shown), as expected from prior reports of patch clamp recordings in rodent striatum brain slices^59^. Conversely, the mTFP1^+^ Pvalb-Pthlh interneurons had a relatively hyperpolarized resting membrane potential, low input resistance, and showed the expected fast-spiking phenotype upon suprathreshold 1s current injection steps (**Figure 7G**). FI (frequency/current) curves were measured from each recorded neuron and used to extract various intrinsic electrophysiological features for comparison. We constructed 3D plots using the most discriminatory sets of electrophysiological features (i.e., FI slope, resting membrane potential, and AP half width, or alternately, FI slope, upstroke downstroke ratio, and input resistance) and observed clear separation of the three distinct interneuron populations into three different spatial domains, consistent with their known signature electrophysiological properties in rodents^26,45^ and confirming the high specificity of viral labeling in the triple RO multiplex experiment (**Figure 7I-J**). PERMANOVA statistical analysis using all five features also showed these cell populations were significantly different (F= 51.071 and p= 0.001) with pairwise PERMANOVA post hoc confirming differences between all groups (p<0.01). This experiment demonstrates that multiple distinct striatal interneuron types can be simultaneously labeled in adult C57Bl6/J mice for detailed analysis of intrinsic properties by patch clamp recording in acute brain slices. It is expected this approach can be applied to study cell type specific synaptic connectivity and neuromodulation in the striatum as well.

### Cross-species conservation of striatal enhancer activity

Although our enhancer AAV screening pipeline was developed for the mouse brain, we note that the validated striatal cell type enhancers were derived from both mouse and human genomic sequence. Additionally, the ability to target specific cell types of the striatum has clear therapeutic potential in humans, but this depends significantly on whether their specificity is conserved across species. We thus investigated both the evolutionary conservation of chromatin accessibility at orthologous enhancer loci in multiple species, as well as an assessment of the conservation of enhancer activity and cell type specificity in the rat and macaque brain.

To explore the evolutionary conservation of striatum-enriched chromatin accessibility at enhancer genomic loci selected as viral tools, we compared previously published bulk ATAC-seq on striatal and cortical tissues from human, mouse, macaque, and rat. Five enhancers derived from either mouse or human genomic sequences were chosen based on their high specificity of labeling in mouse striatum: three pan-MSN enhancers (AiE0441h, AiE0447h, and AiE0367h_C2), one D1 MSN enhancer (AiE0779m), and one D2 MSN enhancer (AiE0452h). Our analysis revealed that in all but one case, the ortholog of AiE0441h in rat, these enhancers exhibited enriched open chromatin activity in striatum compared to cortex across species (**Figure 8A**). This conserved pattern of striatal-enriched chromatin accessibility highlights the evolutionary conservation of regulatory elements associated with these enhancers.

**Figure 8.**
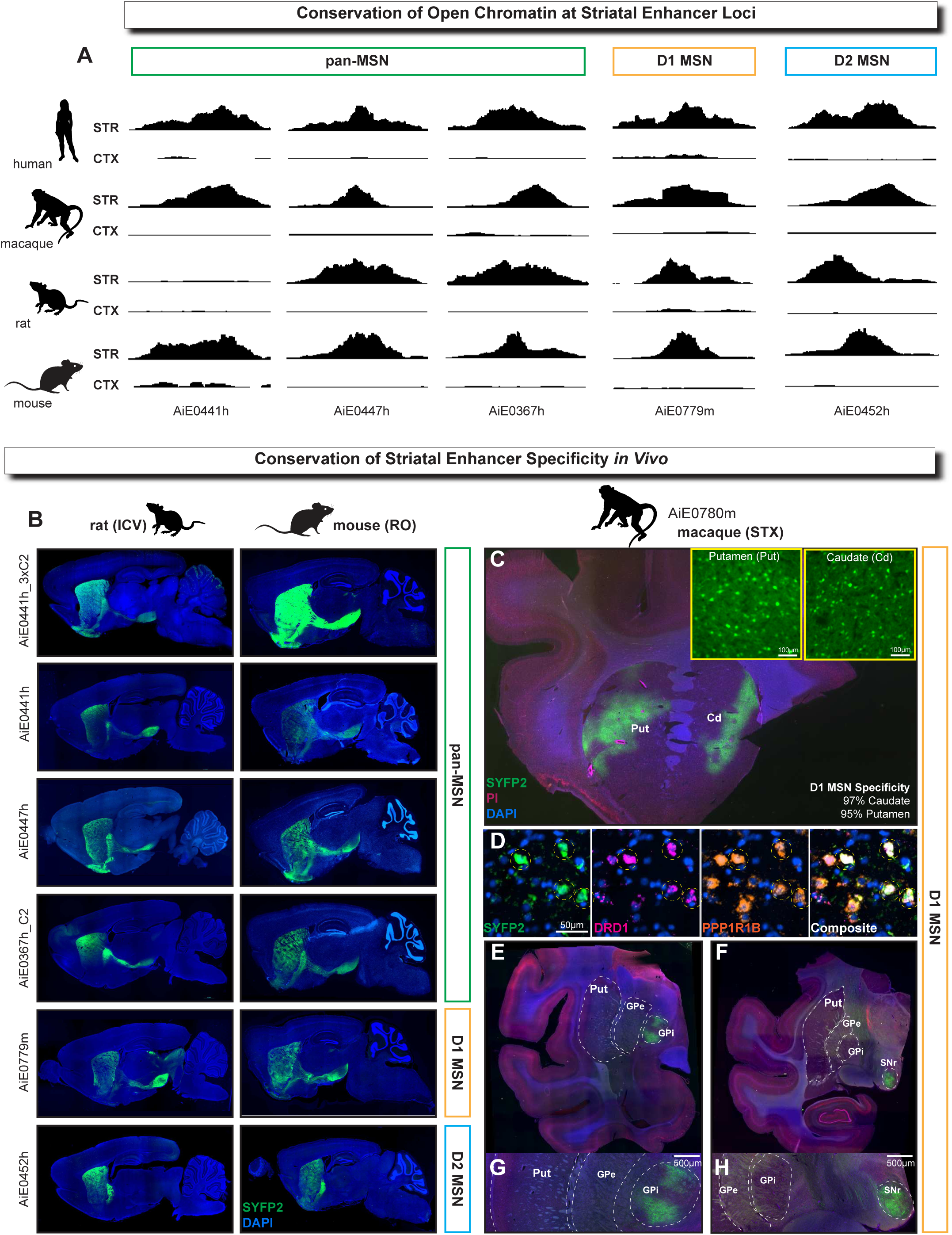
Cross-species conservation of striatal enhancer open chromatin and enhancer AAV activity *in vivo*. A) Assessment of evolutionary conservation of open chromatin activity. Bulk ATAC-seq analysis showing open chromatin regions in the cortex and striatum of human, macaque, rat, and mouse at conserved orthologs of f ive selected enhancer loci. These enhancers include three pan-MSN enhancers, one D1 MSN enhancer, and one D2 MSN enhancer. ATAC-seq traces demonstrate conserved higher open chromatin activity in the striatum compared to the cortex for all enhancers except the ortholog of AiE0441h in rat, which shows no chromatin accessibility in either cortex or striatum. B-H) Conservation of striatal enhancer specificity *in vivo*. B) Enhancer AAVs corresponding to the set of enhancers shown in (A), including an additional optimized variant (AiE0441h_3xC2), were injected RO into mouse and ICV into rat. Sagittal brain sections show conserved patterns of enhancer activity in mouse versus rat brain for native SYFP2 f luorescence (green), with DAPI counterstain (blue). Images were not acquired under matched conditions and have been adjusted to optimally highlight brain wide expression patterns for enhancer specificity comparison purposes. C-H) Stereotaxic injection of AiE0780m driving SYFP2 (AiP12610) in macaque caudate and putamen using the Brainsight veterinary surgical robot C) Coronal section of striatum injection site with red=propidium iodide, blue=DAPI, green=native SYFP2. Insets of caudate (cd) and putamen (put) show native SYFP2 only. D) Example images of RNAscope from Putamen injection site. Yellow circles indicate SYFP^+^DRD1^+^PPP1R1B^+^ cells. E-H) Coronal sections posterior to the injection site showing axon projections through the GPi and SNr indicative of D1 MSNs with red = propidium iodide, blue=DAPI, green = native SYFP2.

We next evaluated whether these striatal-specific enhancers would show conserved activity in the rat brain *in vivo*. We injected this set of enhancer AAV vectors into P1 rat pups by ICV route and assessed the SYFP2 expression pattern in brain at P19. We found that the brain expression patterns matched the expected pan-MSN, D1 MSN, or D2 MSN activity and were clearly conserved between rat and mouse in all cases (**Figure 8B**). Notably, even in the example where the native rat ortholog of human genomic enhancer AiE0441h showed no chromatin accessibility in the striatum, this enhancer still drove pan-MSN specific reporter expression in the rat brain. We also tested the optimized enhancer AiE0441h_3xC2 and observed stronger SYFP2 expression with more complete MSN coverage throughout the striatum compared to the original full-length enhancer AiE0441h, supporting that enhancer core concatenation is a generalizable strategy for improving enhancer strength while retaining specificity in multiple mammalian species.

Finally, to further explore cross species applications, we performed a series of *in vivo* injections into macaque caudate and putamen. We first performed individual injections of exemplary D1 MSN enhancer AiE0780m driving SYFP2 into macaque caudate and putamen, and D2 MSN enhancer AiE0452h driving mTFP1 into macaque putamen and found dense MSN labeling in the caudate and/or putamen regions in anterior striatum (**Figure 8C-H, S10A-B**). In line with D1 MSN projections, we observed AiE0780m driven SYFP2^+^ axon terminal labeling in the GPi and SNr, with no SYFP2^+^ axon terminal labeling observed in GPe (**Figure 8C-H**). mFISH analysis using RNAscope confirmed >95% D1 MSN specificity across both putamen and caudate injection sites (**Figure 8D**). Furthermore, AIE0452h driven mTFP1 was 97% specific for D2 MSNs in the putamen injection site (**Figure S10B**).

We also performed an *in vivo* duplex injection experiment combining the same D1 MSN and D2 MSN enhancer AAVs to evaluate viability and potential crosstalk between these two distinct enhancers when co-injected in macaque (**Figure S10C-F**). As in the single injection experiments, we found distinct MSN dense labeling throughout the targeted anterior regions of the caudate and putamen. We observed highly enriched SYFP2^+^ axon terminal labeling in the GPi and mTFP1 axon terminal labeling in the GPe (**Figure S10C-F**). While 95% of cells were exclusively positive for either mTFP1 or SYFP2 by IHC analysis, we did detect weak signal from mTFP1^+^ axon terminal labeling in the GPi, but not SYFP2^+^ in the GPe (**Figure S10F**). Since we observed near perfect mutual exclusivity of labeling specificity with duplex mouse RO injection experiments (**Figure 7**), it is possible that the high multiplicity of infection achieved with stereotaxic injection could contribute to enhancer AAV transcriptional crosstalk via the mechanisms previously described^83^. Future work on enhancer AAV multiplexing should focus on how to mitigate this putative enhancer crosstalk under high multiplicity of infection conditions, and other sources of technical noise^61^. Nonetheless, these collective findings underscore the robustness of these enhancers as tools for striatal cell type-specific targeting from rodent to primate brain.

## DISCUSSION

We provide a deeply validated toolbox of enhancer AAV vectors for accessing, monitoring, and perturbing major striatal neuron subclasses *in vivo*. These viral vectors are shown to achieve high specificity of labeling in the striatum following systemic (RO) delivery in mice using multiple independent approaches for molecular validation. We report our best-in-class enhancer AAV vectors for pan-MSN class as well as five major striatal neuron subclasses, each with at least one or more exemplary enhancer achieving ≥90% specificity of SYFP2 reporter expression following RO injection in mice. Additionally, we compared specificity of labeling by three different routes of administration in mice (RO, ICV, and STX injection) and report optimal dose ranges for each route as a helpful guide to future users. This work nicely complements two companion studies comparing how different routes of administration influence enhancer specificity and expression level for predominantly cortical cell type targeting enhancer AAV vectors^59^. Continued rigorous analysis and standardized testing along these lines will help ensure high reliability and reproducibility of labeling with the rapidly growing toolbox of cell type targeting enhancer AAV vectors for brain research.

Importantly, enhancer AAV vectors confer several unique advantages over existing transgenic mouse or rat lines in use to target striatal neuron populations. First, the enhancer AAV vectors can be rapidly modified in a matter of days to insert alternate transgene cargos, as we have demonstrated using contrasting colors of fluorescent proteins (e.g., SYFP2, mTFP1, and tdTomato), Cre or Flp recombinases, optogenetic actuator variants (ChR2 and CoChR^60^), and genetically encoded calcium indicators (jGCaMP8 variants^62^). We demonstrated that exemplary enhancers like our optimized D1 MSN enhancer AAV vector was sufficient to drive meaningful levels of transgenes such as ChR2 to confer robust optogenetic control of neuron firing in acute brain slices and of motor behavior in awake-behaving mice. Second, the enhancer AAV vectors can be injected into any genotype of mice (e.g., wild type or mutant strains such as disease models) at any age without the need for complicated and expensive breeding paradigms or complex genotyping requirements. This approach can save cost and improve efficiency by shortening the time for experiments. Third, the enhancer AAV vectors can readily be packaged with different AAV capsid variants such as AAV-retro^63,64^ to impart new functionality and extend their utility, in this case for axon projection mapping and intersectional circuit tracing. Most importantly, these enhancer AAV vectors can be applied to target and perturb genetically defined homologous cell types across diverse mammalian species beyond the mouse model, which addresses a critical unmet need for advancing studies of brain evolution, comparative cellular neurobiology, and development of cell type-specific therapies for treating human brain disorders.

We focused extensively on developing exemplary enhancers for direct (D1) and indirect (D2) pathway MSNs, as these neuron types have been the topic of intensive investigation in the context of basal ganglia circuit connectivity and neurophysiology^51,65,66^, action selection and movement disorders^9,40,67^ ^70^, neuromodulation and drugs of abuse^71^ ^74^ for many decades. Progress in dissociating the roles of the direct versus indirect pathway striatal MSNs has been greatly accelerated by the development and broad distribution of BAC transgenic mouse lines for expression of EGFP or Cre recombinase^73,74^. We identified and validated two distinct enhancers, AiE0779m and AiE0780m, for targeting the D1 MSN subclass. These enhancers were discovered proximal to known direct pathway MSN marker genes *Slc35d3* and *Pdyn*, respectively. Enhancer core concatenation was successful in increasing strength while preserving specificity for AiE0779m. As such, AiE0779m_3xC2 was initially pursued as the best-in-class D1 MSN enhancer. However, in comparing different routes of administration we found that AAV vectors containing AiE0779m or AiE0779m_3xC2 (regardless of transgene cargo) exhibited loss of specificity following stereotaxic injection in the dorsal striatum, and this loss of specificity could not be circumvented by a simple titration of viral dose. In contrast, AAV vectors with AiE0780m, despite driving weak SYFP2 expression by RO injection route, were able to achieve highly specific and robust D1 MSN expression following stereotaxic injections. This important finding illustrates that some enhancers will be better suited to certain experimental applications more so than others, as opposed to the concept of a single best-in-class enhancer for all applications, and this is a major driver of why we pursued and validated multiple enhancers for each striatal neuron subclass. On a related note, we would like to emphasize that enhancers showing weak but specific activity by RO delivery should not be overlooked for applications requiring direct stereotaxic injection into brain parenchyma. Such enhancers may prove to be exceptional in that context with adequate empirical testing.

Our exemplary D2 MSN enhancer AiE0452h exhibited high specificity of labeling with direct expression of fluorescent protein by all three routes of AAV administration. AiE0452h was also ideal for driving the attenuated iCre mutant (R297T) and FlpO for D2 MSN specific recombination in multiple different Cre- or Flp-dependent reporter mouse lines following RO or stereotaxic injection. In contrast, we observed that RO injection of enhancer AAV vectors nearly always resulted in suboptimal direct expression of GCaMP and ChR2 variants (often with little or no detected expression of the fluorescent reporter gene based on native fluorescence), and this was true even for exemplary enhancers such as AiE0452h. We propose this could be readily improved by development of more efficient capsids for blood brain barrier crossing in mouse brain or stronger minimal promoters compared to the ones we have employed. Until then, stereotaxic injection remains the preferred route for direct enhancer driven expression of transgene cargos for optical monitoring and manipulation of neuronal activity. We also created AiE0452h_3xC2 with stronger expression in D2 MSNs of the striatum but observed unexpected off-target activity in L5 extratelencephalic-projecting neurons of the neocortex. Nonetheless, this boost in striatal D2 MSN expression strength could be leveraged for improved GCaMP or ChR2 expression by direct stereotaxic delivery to the dorsal striatum without spread to the cortex region. The depth of our enhancer AAV toolbox offers many different solutions to achieve the desired transgene expression in specific striatal neuron populations using either direct enhancer driven transgene expression, or with Cre or Flp recombinase mediated expression approaches (and potentially sophisticated Cre <u>and</u> Flp intersectional approaches). Further *in vivo* analysis will be required to fully delineate the relative strengths and weaknesses of these different enhancer AAV expression strategies with different functional transgene cargos and to evaluate any differences in tolerability of expression (short and long term) across the different striatal neuron subclasses. Future efforts to refine striatal cell type taxonomies will also reveal further cell type-specific enhancers, allowing us to target more specific subsets of MSNs, including populations restricted to patch or matrix^50,75^, ventral populations such as those localized to the Islands of Calleja (ICjs) and Neurochemically Unique Domains in the Accumbens and Putamen (D1-NUDAPs)^32,76^ ^78^, as well as so-- and D2-coexpressing populations^29,79^.

Our multiplexing experiments were designed with the fact in mind that many modern neuroscience research labs commonly rely on complex transgenic mouse line crosses (either alone or in combination with viral vectors^80,81^) to generate large cohorts of experimental animals for behavioral, electrophysiological, neurochemical, and anatomical studies to uncover fundamental properties of brain cell types and circuits. The cost can be exorbitant and prohibitive for all but the most well-funded laboratories. For many neuroscience researchers these collections of enhancer AAV vectors can be a welcome alternative and should enable previously challenging or infeasible experimental approaches. For example, we have shown multiplexed labeling of direct versus indirect pathway MSNs together with striatal cholinergic neurons with contrasting fluorescent proteins, as well as triple labeling of striatal Pvalb-Pthlh, Sst-Chodl, and cholinergic interneuron subclasses in wild type C57 mice following a simple RO injection procedure. These validated cell type targeting combinations alone could support wide ranging experiments into striatal cell type and circuit function without the need to invoke any transgenic rodent lines. We have shown direct applicability of striatal cell type targeting enhancer AAVs for use in rats, where availability of transgenic driver and reporter lines is much more limited than for mouse (but see also^82,83^). We envision this approach will also greatly benefit those characterizing the cellular effects of gene knockout mutations or utilizing genetic mouse models of basal ganglia disorders (e.g., mouse models Parkinson’s disease, obsessive-compulsive disorder, and autism spectrum disorders), as well as efforts to use cell type specific AAV vectors for Cre recombination in floxed mouse lines with temporal and spatial precision^84^ and without the risk of confounds like germline recombination^85^ or to achieve cell type specific AAV CRISPR/Cas9^86^ in building on the current capabilities^87,88^. We demonstrated targeted patch clamp recording of three major striatal interneuron subclasses in acute brain slices following triple RO injection in adult mice for analysis of signature neuron intrinsic properties, an approach which can readily be applied in mouse disease models to greatly accelerate efficiency and speed of experimental progress. Direct enhancer driven ChR2 expression can be used for optotagging *in vivo* and reliable identification of striatal neuron types for extracellular recording during behavior as shown in mice^89^ ^92^, but also now extensible to diverse mammalian species.

The enhancer AAV toolbox is not without limitations. Enhancer transcriptional crosstalk has recently been recognized as a major limitation for multiplex screening or targeting using cell type specific enhancer AAV^93^, and as such should be carefully considered in developing experimental designs.

Enhancer crosstalk is a process whereby two or more distinct enhancer AAV genomes occur within the same cell and can interact (likely though concatenation of DNA) to cross-activate spurious transgene expression leading to confounding results, namely loss of intended cell type specificity. While we successfully achieved cases of enhancer multiplexing without overt crosstalk in this study, we also identified specific combinations that did exhibit crosstalk. Our work adds to the intrigue around cell type enhancer crosstalk mechanisms by revealing two important and previously unknown principles. First, we observed that crosstalk between distinct cell type specific enhancers can occur in a brain region specific manner, which we speculate may relate to the effective strength of enhancer driven transgene expression across brain regions. In support of this view, AiE0140_3xC2 drives very strong expression in cortex Pvalb interneuron subclass but weaker expression in striatum Pvalb-Pthlh interneuron subclass. Accordingly, crosstalk was observed in combination with AiE0743m_3xC2 in the cortex but not striatum. A second surprising finding is that crosstalk between cell type enhancer AAV vectors can be mitigated (or modulated) by judicious introduction of select other cell type enhancers, which must be determined through empirical testing. While the underlying detailed molecular mechanisms behind these observations remain elusive, these findings provide important context and clues about cell type enhancer AAV multiplexing in the brain. Until these mechanisms are better understood, it is necessary to test and carefully evaluate enhancer crosstalk in any experimental paradigm requiring multiplexing. In the short term, one at a time and low multiplexing of enhancer AAVs will be the most reliable option with the least potential confounds.

In summary, we present a comprehensive enhancer AAV toolbox that enables precise targeting and manipulation of distinct neuronal subclasses in the striatum. Our collection provides a valuable new resource for the neuroscience community that will facilitate in-depth studies of striatal cell type and circuit function across diverse mammalian species including non-human primates. Given that AAV vectors have been shown to be safe and effective in humans, our collection of striatal cell type specific enhancer AAV vectors may also hold great promise for targeted AAV gene replacement therapies or circuit therapies to treat various disorders of the basal ganglia.

## METHODS

### Animals

All procedures involving mice and rats were approved by the IACUC at the Allen Institute for Brain Science (protocols 2004, 2105, 2301, 2306, 2406, and 2010). Mouse tissue was obtained from 4 to 12- week-old male and female pure C57Bl6/J or Drd1a-tdTomato line 6 hemizygous mice^42,43^ and adult Ai14 and Ai65F heterozygous mice. Animals were provided food and water ad libitum and were maintained on a regular 12-h day/night cycle with no more than five adult animals per cage. Timed-pregnant adult female Sprague-Dawley rats were ordered from Charles River Laboratories and acclimated several days after shipment prior to birth of the pups. Rat pups were tattooed at 1-day postnatal (P1) for identification purposes and used the same day for unilateral intracerebroventricular (ICV) injection of AAV vectors. Rat pups were placed back into the home cage with littermates and the dam until they were euthanized at P19 for brain harvest and transgene expression analysis.

### Enhancer selection and genomics analyses

Marker genes for major mouse striatal neuron classes and subclasses were determined from published studies^9^, databases of transgenic mouse lines, e.g., GENSAT (https://www.gensat.org/^86^), as well as using the Allen Brain Atlas and AGEA tool (https://mouse.brain-map.org/agea^94^) and DropVis mouse scRNA-seq data browser (http://dropviz.org/^35^). To identify short genomic DNA fragments (∼250bp-750bp) representing putative cell type specific striatal enhancers, we analyzed publicly available ATAC-seq datasets including Roussos lab Brain Open Chromatin Atlas (BOCA^36^) human postmortem bulk ATAC-seq dataset (https://labs.icahn.mssm.edu/roussos-lab/boca/), as well as the *Cis*-element Atlas (CATlas^87^) mouse snATAC-seq dataset (http://catlas.org/mousebrain/#!/). We searched for differentially accessible open chromatin peaks within ∼500kb of the top marker gene loci such as *Drd1*/*DRD1*, *Drd2*/*DRD2*, *Adora2A*/*ADORA2A*, *Penk*/*PENK*, *Enk*/*ENK*, *Gpr6*/*GPR6*, etc. Enhancer candidates were prioritized based on strong accessibility in striatal brain regions and/or cell types in comparison to all non-striatal regions and cell types. For analysis of Roussos BOCA the putamen and nucleus accumbens were available as independent ROIs/tracks, thus enabling a search for candidate enhancers with possible enrichment in dorsal versus ventral striatum. Bulk ATAC-seq data was deemed suitable for abundant striatal neuron populations (e.g., pan-MSN and direct versus indirect pathway projecting MSN subclasses) but presumably would not be suitable for rare cell types (e.g., various striatal interneuron subclasses). As such, CATlas mouse snATAC-seq dataset was used to search for enhancer candidates with predicted activity in both rare and abundant striatal neuron populations.

Accessibility counter screening in human whole body was performed by first downloading bigwig files from Zhang et al.^95^ (http://catlas.org/humanenhancer/#!/). To assess the accessibility of each putative enhancer in human tissues we used ‘multiBigwigSummary’ from deeptools. This tool measures for each enhancer the average number of fragments within the enhancers genomic coordinate per tissue. LiftOver coordinates in hg38 were used for all enhancers found in mouse datasets (all enhancers ending in an “m”) except for AiE0769m, which does not have an orthologous enhancer identifiable by liftover in hg38.

### Enhancer cloning into AAV vectors

Enhancers were cloned from human or C57Bl/6J mouse genomic DNA using enhancer-specific primers and Phusion high-fidelity polymerase (M0530S; NEB). Individual enhancers were then inserted into a recombinant single-stranded pAAV backbone that contained the beta-globin minimal promoter, fluorescent reporter gene (typically SYFP2), minimal woodchuck hepatitis posttranscriptional regulatory element (WPRE3), and bovine growth hormone polyA using standard molecular cloning as previously described^39^. Plasmid integrity was verified via Sanger DNA sequencing and in some cases restriction digestion with agarose gel electrophoresis to confirm intact inverted terminal repeats (ITRs). In some cases, synthetic enhancer sequences such as concatenated cores were gene synthesized and subcloned into AAV vectors using standard restriction enzyme digestion and ligation. All AAV plasmids were propagated in NEB stable *E.coli* at 30°C growth condition to prevent spurious DNA rearrangements.

### AAV packaging and titer determination

Small-scale crude AAV preps were generated by triple transfecting 15 µg ITR plasmid, 15 ug AAV capsid plasmid, and 30 ug pHelper (Cell Biolabs) into one 15-cm plate of confluent HEK-293T cells using PEI Max (Polysciences Inc., catalog # 24765-1). At one day post-transfection we changed the medium to low serum (1% FBS), and after 3 days the cells and supernatant were collected, freeze-thawed 3x to release AAV particles, treated with benzonase nuclease (MilliporeSigma catalog # E8263-25KU) for 1 hr to degrade free DNA, then clarified (3000g x 10min) and concentrated to approximately 150 µL by using an Amicon Ultra-15 centrifugal filter unit (NMWL 100 kDa, Sigma #Z740210-24EA) at 5000g for 30-60 min, yielding a titer of approximately 1.0E+13 to 1.0E+14 vg/mL. An efficient and cost-effective protocol for small scale crude AAV preparation has been shared on protocols.io (https://www.protocols.io/view/production-of-crude-aav-virus-extract-yxmvmx7r9l3p/v6) by the Allen Institute team. This protocol typically yields 5E+12vg from one 15 cm plate of AAV293 cells using the conventional adherent cell culture and triple transfection method with serotype PHP.eB.

For large-scale gradient preps, we transfected 10 x 15-cm plates of cells and purified by iodixanol gradient centrifugation. For measuring virus titers, we used ddPCR (Bio Rad; QX 200 Droplet Digital PCR System). We used primers against AAV2 ITR for amplification. Seven serial dilutions with the factor of 10 ranging from 2.5×10^-2^ to 2.5×10^-8^ were used for the measurement. Serial dilutions of 2.5×10^-5^ to 2.5×10^-8^ were used for fitting the dynamic linear range. Viral titer was calculated by averaging virus concentration of two dilutions within the dynamic linear range. A positive control of a known viral titer, and a negative control with no virus was also run along with all the samples.

### Retro-orbital (RO), intracerebroventricular (ICV) and stereotaxic (STX) injection delivery of AAV vectors

Adult C57Bl6/J mice were briefly anesthetized by isoflurane anesthesia and injected with crude PHP.eB^88^ serotyped AAV virus at a dose range of 1E+10 to 1E+12vg into the retroorbital sinus of one eye in a maximum volume of 100µL. Virus stocks were diluted in sterile 1X phosphate buffered saline (1XPBS) solution as needed to achieve the intended dose and volume. For initial screening with enhancer AAVs driving SYFP2, we routinely used 5E+11vg as the standardized RO dose. For ICV injections, C57BL/6J or CD-1_IGS mouse pups and Sprague Dawley rat pups (P0-P2) were injected with 3E+10 to 1.5E+11vg of concentrated AAV vector into the lateral ventricle(s). In most experiments animals received unilateral ICV injection but in a subset of experiments some mice received bilateral ICV injections to evaluate and compare brain wide viral transduction and transgene expression. For mouse *in vivo* optogenetics experiments, C57Bl6/J mice received a unilateral (left or right) injection of 3E+10vg concentrated AAV vector stock by ICV into the lateral ventricle at P2. For stereotaxic injection surgery, C57Bl/6J, Ai14 het, Ai65F het, and Drd1-Cre adult mice were deeply anesthetized to a surgical plane using an isoflurane vaporizer and placed into the stereotaxic injection frame. AAV virus was injected bilaterally into the dorsal striatum region (dSTR) using the following coordinates (in mm) relative to Bregma: anterior/posterior (A/P) 0.8, medial/lateral (M/L) ±1.6 to 2.0, and dorsal/ventral (D/V) 2.6 to 3.0. A total volume of 500 nL containing 1E+12 to 1E+13 vg/mL virus stock was delivered at a rate of 50 nL per pulse with a Nanoject II pressure injection system. Before incision, the animal was injected with Bupivacaine (2-6 mg/kg) and post injection, the animal was injected with ketofen (2-5 L) to provide analgesia. Mice that underwent STX injections were euthanized after 3-5 weeks post injection, transcardially perfused with 1XPBS followed by 4% paraformaldehyde (PFA), and the brains were dissected for further analysis. In a subset of STX injection experiments for acute brain slice electrophysiology, mature adult mice were sacrificed for photocurrent measurement experiments at 1-2 months post injection.

### Tissue processing for slide-based epifluorescence imaging

Mice were anaesthetized with isoflurane and perfused transcardially with 10 m L of 0.9% saline, followed by 50 mL of 4% PFA. The brain was removed, bisected along the midsagittal plane, placed in 4% PFA overnight and subsequently moved to a 30% sucrose solution until sectioning. From the left hemisphere, 30μm sections were ontained along the entire mediolateral axis using a freezing, sliding microtome. Five sagittal ROIs, roughly 0.5, 1, 1.5, 2.3 and 3.5 mm from the midline, were collected and stained with DAPI ((4’,6-diamidino-2-phenylindole dihydrochloride) and/or propidium iodide (PI) to label nuclei and to reveal cellular profiles, respectively. Stained tissue sections were slide mounted using Vectashield hardset mounting medium (Vector Laboratories, catalog # H-1400-10) and allowed to dry for 24 hours protected from light. Once the mounting medium hardened, the slides were scanned with Aperio VERSA Brightfield epifluorescence microscope (Leica) in the UV, green, and red channels, illuminated with a metal halide lamp. After passing QC, digitized images were analyzed by manual scoring of virus-mediated fluorescent protein expression throughout the brain with emphasis on striatal regions including caudoputamen, nucleus accumbens, and olfactory tubercle. Images were not acquired under matched conditions and have been adjusted to optimally highlight brain wide expression patterns for enhancer specificity comparison purposes.

### Serial two-photon tomography (Tissuecyte)

Mice were perfused with 4% PFA. Brains were dissected and post-fixed in 4% PFA at room temperature for 3 6 h and then overnight at 4 °C. Brains were then rinsed briefly with PBS and stored in PBS with 0.01% sodium azide before proceeding to the next step. Agarose was used to embed the brain in a semisolid matrix for serial imaging. After removing residual moisture on the surface with a Kimwipe, the brain was placed in a 4.5% oxidized agarose solution made by stirring 10 mM NaIO4 in agarose, transferred through phosphate buffer and embedded in a grid-lined embedding mold to standardize its placement in an aligned coordinate space. The agarose block was then left at room temperature for 20 min to allow solidification. Brain tissue was additionally supported by generating a polyacrylamide network throughout the agarose block and spanning the brain-agarose interface. The agarose block was first left at 4°C overnight in a solution of 4.5% Surecast (Acrylamide:Bis-acrylamide ratio of 29:1) with 0.5% VA-044 activator, diluted in PBS. Agarose blocks were then placed back into new embedding molds containing a small amount of acrylamide solution and the top surface covered with parafilm (to reduce exposure to oxygen). Finally, specimens are baked for 2 hours at 40°C and then stored in PBS with 0.1% sodium azide at 4°C until ready to image. The agarose block was then mounted on a 1 × 3 glass slide using Loctite 404 glue and prepared immediately for serial imaging.

Image acquisition was accomplished through serial two-photon (STP) tomography using six TissueCyte 1000 systems (TissueVision, Cambridge, MA) coupled with Mai Tai HP DeepSee lasers (Spectra Physics, Santa Clara, CA). The mounted specimen was fixed through a magnet to the metal plate in the center of the cutting bath filled with degassed, room-temperature PBS with 0.1% sodium azide. A new blade was used for each brain on the vibratome and aligned to be parallel to the dorsoventral axis.

Brains were imaged from the caudal end. We optimized the imaging conditions for both high-throughput data acquisition and detection of single axon fibers throughout the brain with high resolution and maximal sensitivity. The specimen was illuminated with 925 nm (EGFP, tdTomato, dTomato data) or 970 nm (SYFP2 data) wavelength light through a Zeiss ×20 water immersion objective (NA = 1.0), with 250 mW light power at objective. The two-photon images for red, green and blue channels were taken at 75μm below the cutting surface. This depth was found optimal as it is deep enough to avoid any major groove on the cutting surface caused by vibratome sectioning but shallow enough to retain sufficient photons for high contrast images. In order to scan a full tissue section, individual tile images were acquired, and the entire stage was moved between each tile. After an entire section was imaged, the x and y stages moved the specimen to the vibratome, which cut a 100-m section, and specimen to the objective for imaging of the next plane. The blade vibrated at 60 Hz and the stage moved towards the blade at 0.5 mm per sec during cutting. Images from 140 sections were collected to cover the full range of mouse brain. It takes about 18.5 h to image a brain at an x,y resolution of ∼ 0.35 m per pixel, amounting to ∼750 GB worth of imaging, sections were retrieved from the cutting bath and stored in PBS with 0.1% sodium azide at 4°C.

### Barcoded enhancer AAV multiplex assay

#### Cloning and synthesis of barcoded enhancer AAVs

The 8bp barcode strategy was derived from Guo et al.^96^ with all fourteen barcodes used in this study designed to have a Hamming Distance >2. Barcodes were ordered as single-stranded forward and reverse compliment oligonucleotides from Integrated DNA Technologies (IDT) with 20bp overlapping homology regions on either side of the XhoI site into each parent vector. Each oligonucleotide stock was resuspended in water to 100uM and then combined and diluted 1:500 with its pair. Oligo pairs were annealed by boiling at 100C for 5 minutes and slowly cooling to room temperature over 30 minutes. The annealed oligonucleotides were cloned into the parent vectors using In-Fusion HD Cloning Kit (Takara 639650) and transformed into chemically competent Stbl3 E. coli (ThermoFisher C737303). Individual colonies were selected for on 100 ug/mL carbenicillin plates. Plasmids were maxiprepped (Qiagen 12162) and sequence-verified using Azenta Sanger sequencing. Completed plasmids were individually packaged into PHP.eB capsid using the small-scale crude prep method (please reference above section: AAV packaging and titer determination).

#### Validation of barcoded enhancer AAVs

To confirm 3 UTR barcodes did not affect enhancer expression levels, barcoded enhancer AAVs were injected individually into mice and compared using epifluorescence imaging to original constructs using the same dose (5E+11 vg/mouse) and route of administration (RO).

#### Single cell isolation of tissue from mice injected with barcoded enhancer AAVs

Barcoded enhancer AAVs were pooled for a total of 9.8E11 vg (7E10 vg per enhancer AAV) and diluted in sterile 1xPBS for a final volume of 100uL. Pooled AAVs were injected retro-orbitally into adult mice. After 4 weeks incubation, single cells were isolated as described previously^30^. Mice were anesthetized with Avertin and perfused transcardially with ice-cold Cutting Buffer containing 110mM NaCl, 2.5mM KCl, 10mM HEPES, 7.5mM MgCl2, 25mM glucose, and 75mM sucrose. Following perfusions, brains were extracted and placed in a petri dish containing Cutting Buffer to generate 2mm coronal slices using a stainless steel brain matrix (Stoelting 51386). Slices containing dorsal striatum were then transferred to a second petri dish with Dissociation Buffer containing 82mM Na2SO4, 30mM K2SO4, 10mM HEPES, 10mM Glucose, and 5mM MgCl2 and dorsal striatum was isolated using a disposable 2mm tissue punch (VWR 95039-098). Tissue pieces were then placed immediately in 10mL conical tubes containing 5mL of room temperature Enzyme Buffer containing 3mg/mL Protease XXIII (Sigma-Aldrich P5380) and 10 units/mL of Papain (Worthington LK003150) dissolved in Dissociation Buffer.

Submerged slides were immediately transferred to a 34C water bath for 1 hour. Following digestion, conical tubes were placed directly on ice and the supernatant was carefully removed and replaced with 10mL Stop Solution containing 1mg/mL Trypsin Inhibitor (Sigma-Aldrich T6522), 2mg/mL BSA (Sigma-Aldrich A2153), and 1mg/mL Ovomucoid Protease Inhibitor (Worthington LK003150) dissolved in Dissociation Buffer. Tissue chunks were triturated using fire-polished glass Pasteur pipets with successively smaller bore holes while avoiding bubbles until a homogenous solution was obtained.

Triturated tissue was centrifuged at 300xg for 10min at 4C, supernatant decanted, pellet resuspended in 5mL of Stop Solution, and then centrifuged again at 300xg for 10min at 4C. Following centrifugation, the supernatant was decanted once more and the cell pellet was resuspended in 500uL-1mL of Dissociation Buffer with 0.1% BSA, 1:500 DAPI (ThermoFisher 62248), and 1:500 Vibrant DyeCycle Ruby Stain (ThermoFisher V10309). The resulting single cell suspensions were filtered using a pre-wet 70µm filter (Miltenyi Biotec 130-098-462) and incubated in the dark at 4C for 30 minutes prior to flow cytometry.

Cell suspensions were sorted for SYFP2 expression using a 130µm nozzle on a BD FACSAria III at a flow rate of <2000 cells per second on purity mode. Live cells were selected for by a DAPI-negative, Ruby-positive gate. Approximately 45,000 SYFP2^+^ and SYFP2^-^ cells were collected in chilled 5mL FACS tubes containing 300uL Dissociation Buffer with 0.1% BSA. The sorted cells were centrifuged at 300xg for 10min at 4C, supernatant decanted, and the pellet resuspended in 80-100uL of Dissociation Buffer with 0.1% BSA on ice. Cell concentrations were estimated using a disposable hemocytometer (Thomas Scientific 1190G82) and 15,336 SYFP2^+^ whole cells were immediately loaded on the 10X v3.1 chip for insertion into the 10X Genomics Chromium controller.

*10X Genomics, custom AAV barcode library generation, and sequencing* transcription, cDNA amplification and library construction were followed. Libraries were sequenced on the Illumina NovaSeq 6000 with a target read depth of 125,000 reads per cell. To generate custom libraries for the AAV barcoded transcripts, 10ng of the remaining amplified cDNA from each sample was first used in a 100uL reaction containing 50uL KAPA Hifi Mastermix (Roche 7958935001), 0.5uL 100uM forward primer that binds to the WPRE3 in the AAV vector 5’ AACTCATCGCCGCCTGCCTTG primer that binds to Read 1 (part of the oligo attached to the 10X gel bead) CTACACGACGCTCTTCCGATCT conditions: 98C for 5 minutes initial denaturation, then 8x cycles of 95C for 15 seconds, 60C for 30 seconds, 72C for 20 seconds, and lastly 72C for 30 seconds. The reaction was purified using 0.7x and 0.9x SPRI beads (Beckman Coulter B23318) and then resuspended in 40uL Buffer EB (Qiagen 19086). A second reaction containing 20uL of product from the first reaction, 0.5uL of 100uM nested forward AATGATACGGCGACCACCGAGATCTACAC**NNNNNNNNNN**ACACTCTTTCCCTACACGACGCTCTT CCGATCT CAAGCAGAAGACGGCATACGAGAT**NNNNNNNNNN**GTGACTGGAGTTCAGACGTGTGCTCTTCCG ATCTACTGACAATTCCGTGGCTCG adapters and indexes to the amplicons. The thermocycler conditions were kept the same as the first PCR except for the cycle number was increased to 12x. The reaction was purified using 0.7x and 0.9x SPRI beads, resuspended in 25uL Buffer EB, and checked for purity and concentration using the High Sensitivity Kit (Agilent 067-4626) on a Bioanalyzer. Custom libraries were pooled prior to sequencing on the Illumina NextSeq 2000 with a target read depth of 40,000 reads per cell.

#### Sequencing, QC filtering, and enhancer specificity calculations

A custom reference was created using the 10x Genomics Cellranger mkref to add the AAV vector sequences to the mouse reference transcriptome (M21, GRCm38.p6). Fastq files were aligned to the custom reference using the 10x Genomics CellRanger pipeline (version 6.1.1) while also including a feature barcode reference with directories to the AAV barcode libraries, as instructed by 10x Genomics Feature Barcode Technol High-quality cells were selected for using stringent QC filtering as described previously^89^. Cells with total reads that contained transcripts from >20% mitochondrial genes, <2000 genes detected, and a doublet score of >0.3 (DouletFinder algorithm from scrattch.hicat https://github.com/AllenInstitute/scrattch.hicat) were removed from the dataset resulting in 8,671 high quality cells. Filtered gene x cell count matrices were then normalized using the logCPM function (also part of the scrattch.hicat R package) and then mapped using the correlation mapping strategy from scrattch.mapping (https://github.com/AllenInstitute/scrattch.mapping) onto the 10X Whole Mouse Brain taxonomy (CCN20230722). Cells that contained <2 AAV transcript UMIs were also discarded. The resulting 6,577 cells were assigned to each enhancer by the expressed enhancer-paired barcode, with 43% (2878/6577) of the cells assigned to more than one enhancer-barcode. We compared the rank order of enhancers from cells with a single barcode detected to cells with >1 barcodes detected and found a highly significant agreement in rank order of enhancers (Kendall’s rank correlation test, T = 0.909, p <0.001). Thus, barcode counts were included from both single and multiple infections. Enhancer specificity was determined by number of cells in each cell subclass with viral vector derived transcripts and enhancer AAV strength was determined by median transcript UMI per cell.

#### Immunohistochemistry

Brain slices were fixed in 4% PFA in phosphate buffered saline (PBS) at 4 °C overnight or up to 48 hours and then transferred to 1XPBS with 0.01% sodium azide as a preservative. Fixed slices were thoroughly washed with PBS to remove residual fixative, then blocked for 1 hr at room temperature in 1XPBS containing 5% normal goat serum and 0.2% Triton-X 100. After blocking, slices were incubated overnight at 4 °C in blocking buffer containing one or more of the following primary antibodies: chicken anti-GFP (Aves, 1:1000 or 1:2000), rabbit anti-red fluorescent protein (Rockland, 1:500), mouse anti-ChAT IgG2b (Atlas Labs, 1:1000), Rabbit anti-nNos (Immunostar, 1:1000), Rabbit anti-Pvalb (Swant, 1:2500). Following the overnight incubation, slices were washed for 15 min three times with 1XPBS and then incubated for 2 hr at room temperature in dye-conjugated secondary antibodies (1:1000; Invitrogen, Grand Island, NY) including BV480 (goat anti-rat)Alexa Fluor 488 (goat anti-chicken), Alexa Fluor 555 (goat anti-rabbit and goat anti-mouse), and Alexa Fluor 647 (goat anti-mouse). Slices were washed for 15 min three times with 1XPBS, followed by 5ug/mL DAPI nuclear staining for 15 min. The slices were then dried on glass microscope slides and mounted with Fluoromount-G (SouthernBiotech, Birmingham, AL). Slides were stored at room temperature in the dark prior to imaging. Whole slice montage images were acquired with NIS-Elements imaging software on a Nikon Eclipse Ti or Ti2 Inverted Microscope System equipped with a motorized stage and epifluorescence illumination with standard DAPI, FITC, TRITC, Cy3, and Cy5 excitation/emission filter cubes. Confocal z-stack images were acquired on an Olympus Fluoview 3000 laser scanning confocal microscope equipped with 488 nm, 543 nm, 594 nm, and 647 nm excitation laser lines.

Specificity of enhancer activity for the target striatal cell subclass or type was quantified as reporter and marker Ab double positive neuron count divided by total reporter Ab positive neuron count and multiplied by 100 to obtain a percentage. The main ROI for analysis was in the center of the dorsal striatum region. A minimum of 50 neurons were counted in each ROI. Completeness of labeling of a given target striatal cell subclass or type was quantified as reporter and marker Ab double positive neuron count divided by total marker positive neuron count and multiplied by 100 to obtain a percentage.

### RNAscope

#### Mouse

We performed RNA FISH using RNAscope Multiplex Fluorescent v1 and v2 kits (Advanced Cell prepared from fresh frozen brains and embedded in optimum cutting temperature compound (OCT; Tissue-Tek). For the v2 kit, 30µm sections were prepared from mice perfused with 4% PFA, the brain extracted and sunk in 30% sucrose before OCT embedding. All coronal sections for both kits were cut using a cryostat (CM3050 S) and mounted on SuperFrost slides (ThermoFisher Scientific). Slides were v2 kit, during the sample preparation and pretreatment step, we increased the slide baking time from 30 minutes in the manual to 1 hour to improve tissue adherence. When developing HRP signal, the HRP blocking step was increased from 15 minutes in the manual to 30 minutes to minimize cross amplification between the different channels/probes. The following probes were used: ChAT (ACD Cat#408731-C2) to label PAL-STR Gaba-Chol cells, Ppp1r1b (ACD Cat#405901-C2 or ACD Cat#405901-C3) to label all MSNs, Drd1 (ACD Cat#461901-C3) to label D1 MSNs, Drd2 (ACD Cat#406501-C3) to label D2 MSNs and SYFP2 mRNA (ACD Cat#590291-C1) for detecting enhancer-driven transcripts. To visualize the probes, we used the following TSA dyes and concentrations from ACD: TSA Vivid 520 (1:1500, Cat# 7534) for SYFP2, TSA Vivid 570 (1:2000, Cat#7535) for ChAT, Drd1, and Drd2, and TSA Vivid 650 (1:5000, Cat#7536) for Ppp1r1b. DAPI (Cat#323108) labelled nuclei. We imaged slides at 40X on a confocal microscope (Leica SP8). Specificity measurements for the dorsal striatum were determined using QuPath cell detection and object classifiers^97^. The DAPI nuclear signal was used to segment cells in images. We did note some bleed-through of probes between the channels (especially with the Ppp1r1b probe which had an exceptionally strong signal). To quantify real signal over background, we trained QuPath object classifiers separately for each probe using individual training images before applying to the ROI. Specificity of enhancer AAVs was calculated as follows: SYFP2^+^Chat^+^ / SYFP2^+^ for cholinergic interneuron enhancers, SYFP2^+^Ppp1r1b^+^ / SYFP2+ for pan-MSN enhancers, SYFP2^+^Drd1^+^ / SYFP2^+^ for D1 MSN enhancers, and SYFP2^+^Drd2^+^ / SYFP2^+^ for D2 MSN enhancers. For AiE0779m enhancer versions, completeness of labeling and SYFP2 intensity was also calculated using QuPath. To normalize for differences in background levels, intensity values per cell were calculated as SYFP2 intensity (a.u.) minus average SYFP2 intensity across SYFP2 negative cells. Completeness was calculated as SYFP2^+^Drd1^+^/Drd1^+^.

#### Macaque

The RNAscope Multiplex Fluorescent v2 kit was used in tandem with the RNA-Protein Co-detection Ancillary kit (ACD) with a few modifications. The tissue was cut using a freezing sliding microtome and mounted SuperFrost slides and used within one month. During sample preparation and pretreatment, we performed the hydrogen peroxide wash first before allowing the sections to air dry for 2 hours on the slide. The slide baking step was increased from 30 minutes to 1 hour for improved tissue adherence. We added an additional 15-minute baking step after the EtOH washes also for improved tissue adherence. Due to loss of native SYFP2 and mTFP1 signal from the v2 kit protocol, we incorporated the RNA-Protein Co-detection kit to perform IHC to visualize the enhancer driven SYFP2 and mTFP1 while using the probes for the marker gene mRNA. We followed the co-detection kit protocol and placed the tissue in primary antibody solution at 4C after the antigen retrieval step where it incubated overnight. The primary antibodies used to visualize the SYFP2 and mTFP1 were chicken anti GFP (1:250, Aves Lab Cat# GFP1020) and rat anti mTFP (1:250, CancerTools Cat#155264). The secondary antibody incubation occurs after the final HRP blocking step. The tissue was placed in secondary antibody solution for 3 hours at room temperature. The secondary antibodies used were goat anti chicken Alexa Fluor 488 (1:500, Invitrogen Cat#A11039) and goat anti rat Alexa Fluor 647 (1:500, Invitrogen Cat#A21247). The probes used were: Drd1 (ACD Cat# 1075621-C2) to label D1 MSNs, Drd2 (ACD Cat#549031-C2) to label D2 MSNs, and Ppp1r1b (ACD Cat# 1075651-C1) to label all MSNs. To visualize the probes, we used the following TSA dyes and concentrations from ACD: TSA Vivid 570 (1:1500, Cat#7535) for Ppp1r1b and TSA Vivid 650 (1:1500, Cat#7536) for Drd1 and Drd2. DAPI (Cat#323108) labelled nuclei. All slides are mounted with ProLong Gold Antifade mounting media (ThermoFisher Scientific Cat#P36930) and allowed to dry overnight before imaging on a Nikon Eclipse Ti2 Inverted Microscope System equipped with a motorized stage and epifluorescence illumination with standard DAPI, FITC, TRITC, Cy3, and Cy5 excitation/emission filter cubes. To quantify specificity, images were loaded into FIJI where brightness/contrast was manually adjusted to eliminate background signal in all channels. Cells were then marked using the Cell Counter plugin based on positivity for GFP, Drd1 or Drd2, and Ppp1r1b. Specificity was calculated as GFP^+^Drd1^+^Ppp1r1b^+^ / GFP^+^ and mTFP^+^Drd2^+^Ppp1r1b^+^ / mTFP^+^.

### SMART-seq v4 sample preparation and analysis (scRNA-seq)

Single cell suspensions from enhancer AAV RO injected mice were prepared for flow cytometry and single cell RNA-seq from brain tissue as previously described^98^. Briefly, for flow cytometry, we perfused mice transcardially under anesthesia with ACSF.1. We harvested the brains, embedded in 2% agarose in PBS, then sliced thick 350µm sections using a compresstome with blockface imaging, then picked the sections containing the dorsal striatum and dissected it out. We then treated dissected tissues with 30U/mL papain (Worthington LK003176) in ACSF.1 containing 30% trehalose (ACSF.1T) in a dry oven at 35°C for 30 minutes. After papain treatment we quenched digestion with ACSF.1T containing 0.2% BSA, triturated sequentially using fire-polished glass pipettes with 600, 300, and 150 micron bores, filtered the released cell suspensions into ACSF.1T containing 1% BSA, centrifuged cells at 100g for 10 min, then resuspended cytometry and sorting on a FACSAria III (Becton-Dickinson). Sample preparation for SMART-Seq was performed using the SMART-Seq v4 kit (Takara Cat#634894) as described previously^98^. Single cells were sorted into 8-well strips containing SMART--Seq reagents were used for reverse transcription and cDNA amplification. Samples were tagmented and indexed using a NexteraXT DNA Library Preparation kit (Illumina Cat#FC-131-1096) with NexteraXT Index Kit V2 Set A (Illumina Cat#FC-131-2001) according to manufacturer’s decreases in volumes of all reagents, including cDNA, to 0.4 x recommended volume. Full documentation for the scRNA-seq procedure Institute data portal at http://celltypes.brain-map.org/. Samples were sequenced on an Illumina HiSeq 2500 as 50 bp paired end reads. Reads were aligned to GRCm38 (genecode v23) using STAR v2.7.1 with the parameter “twopassMode,” and exonic read counts were quantified using the GenomicRanges package for R as described in Tasic et al. (2018). To determine the corresponding cell type for each scRNA-seq dataset, we utilized MapMyCells hierarchical mapping from the Allen Institute http://celltypes.brain-map.org/ according to the posted instructions against the 10x Whole mouse brain taxonomy (CCN20230722). We applied quality control steps to exclude cells from the dataset if they had less than 1,000 genes detected (196 cells; 8%) and if they clearly did not map to basal ganglia cell subclasses corresponding to the dissected brain regions (133 cells; 5%). The remaining 2,178 high-quality cells (87%) were used to determine enhancer AAV labeling specificities across multiple experiments. Below is a list of taxonomy cell classes and subclasses and common names they correspond to:

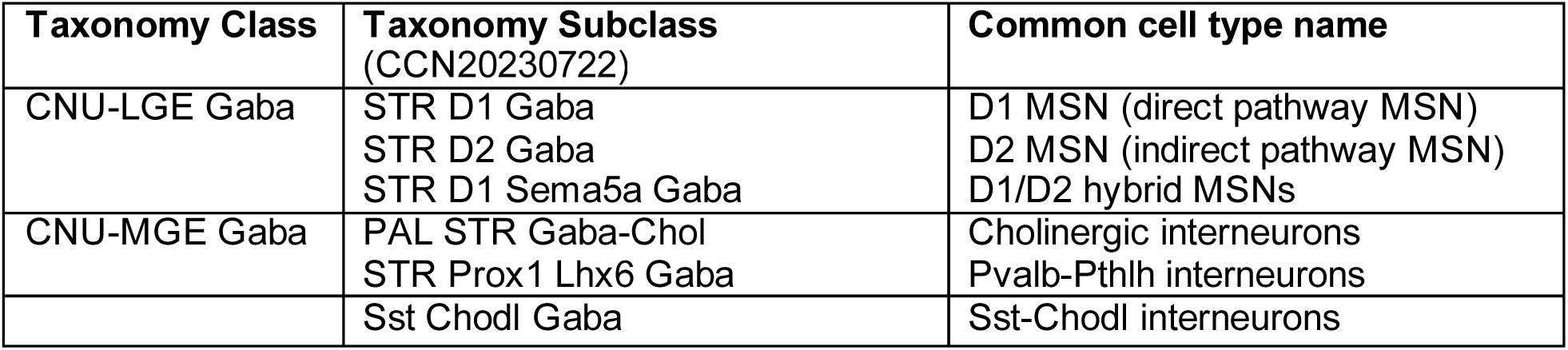

| Taxonomy Class | Taxonomy Subclass (CCN20230722) | Common cell type name |
| --- | --- | --- |
| CNU-LGE Gaba | STR D1 Gaba<br>STR D2 Gaba<br>STR D1 Sema5a Gaba | D1 MSN (direct pathway MSN)<br>D2 MSN (indirect pathway MSN)<br>D1/D2 hybrid MSNs |
| CNU-MGE Gaba | PAL STR Gaba-Chol<br>STR Prox1 Lhx6 Gaba | Cholinergic interneurons<br>Pvalb-Pthlh interneurons |
|  | Sst Chodl Gaba | Sst-Chodl interneurons |

### Analysis of endogenous marker gene expression in enhancer AAV labeled cells

Using the cell x gene matrix from our RO SYFP2+ reporter SSv4 datasets, log2(CPM+1) normalized mRNA transcript counts for select marker genes were first averaged across cells for each enhancer. Enhancers were next grouped by their cell type target and then by whether the enhancer was found proximal or non-proximal to each marker gene. The average of each proximal and non-proximal group for each target cell type was then plotted to assess differences in marker gene expression. The following enhancers (with proximal marker gene they were found near in parentheses) were included in the analysis:

- Pan-MSN: AiE0441h (*Ppp1r1b*), AiE0441h_3xC2 (*Ppp1r1b*), AiE0621h (*Pde7b*)
- D1 MSN: AiE0779m (*Slc35d3*), AiE0779m_3xC2 (*Slc35d3*), AiE0779m_6xC2 (*Slc35d3*), AiE0616h (*Pdyn*), AiE0780m (*Pdyn*)
- D2 MSN: AiE0452h (*Gpr6*), AiE0452h_3xC2 (*Gpr6*), AiE0444h (*Drd2*), AiE0617h (*Adora2a*)
- Pvalb-Pthlh: AIE0140h_3xC2 (*Nrf1*), AiE0951h (*Drd3*), AiE1419m (*Pthlh*), AiE1421m (*Igfpb4*) Sst-Chodl: AiE0600m (*Nos1*), AiE0682h (*Nos1*), AIE0769m (*Pnp2*), AiE1426m (*Nxph2*)
- Cholinergic: AiE0743m (*Tacr3*), AiE0743m_3xC2 (*Tacr3*), AiE0873m (*Slc5a7*), AiE0873m_3xC2 (*Slc5a7*), hChATp (*Chat*)

### Comparative ATAC-seq analyses

Previously published bulk ATAC-seq data from cortex and striatum of human (GSE96949), macaque (GSE159815), Database.

We combined all available replicates available in bigWig format in human (n = 9 primary motor cortex; n = 10 putamen), macaque (n = 2 primary motor cortex, n = 2 caudate), mouse (n = 2 motor cortex; n = 2 striatum), and rat (n = 2 motor cortex, n = 2 striatum) using bigWigMerge with default parameters.

Combined bigwig files were examined using the Integrative Genomics Viewer (IGV), indexing using IGV’s to TDF tool and group autoscaling tracks to ensure comparability of cortex and striatum open chromatin activity within species. Orthologous loci of either mouse- or human-derived enhancers were each o comparative display we centered a 500 bp window around each peak, reversing sequences as needed to preserve the syntenic orientation relative to its closest gene observed in the parent sequence.

### Stereotaxic implantation of optical fibers

Stereotaxic surgeries were performed in accordance with protocols approved by the Stanford Institutional Animal Care and Use Committee. Mice were maintained on a 12/12 light/dark cycle and fed ad libitum. All surgeries were carried out in aseptic conditions while mice were anaesthetized with isoflurane (5% for induction, 0.5%1.5% afterward) in a manual stereotactic frame (Kopf). Buprenorphine HCl (0.1 mg kg-1, intraperitoneal injection) and Ketoprofen (5 mg kg-1, subcutaneous injection) were used for postoperative analgesia. Mice were allowed to recover for at least 1 week before experiments. 200 mm fiber optic ferrules with 0.48 NA were implanted in the same surgery at coordinates +0.5mm AP, +/-1.5mm ML, −3.0mm DV.

### Optogenetic stimulation

Mice were implanted unilaterally with fiber optic ferrules in dorsal striatum (injection coordinates: +/- 1.5 ML; +0.5 AP; −3.0 DV). Mice were tethered to flexible optical fibers connected to a fiber optic commutator (Doric Lenses). Optogenetic light stimulation was delivered with 10 s pulses of 450 mW s prior to light 15 minutes. Equipment for LED light generation, optical fibers, and commutation were from Doric Lenses. The opsin variant ChR2(CRC)-EYFP contains the mutations L132C, H134R, and T159C and was selected due to relative step like activation kinetics with minimal to no desensitization. This enables neurons to exhibit more reliable sustained action potential firing in response to prolonged blue light stimulation relative to ChR2(H134R)-EYFP.

### Tracking of locomotion in optogenetic stimulation

Mice were placed in a darkened, circular open arena 30 cm in diameter with a transparent floor illuminated from underneath with infrared LEDs. Video was acquired with a camera mounted underneath at 65 Hz. DeepLabCut was used to identify the nose and tail base of each mouse^99^. The average of these points was used to identify the center point for quantification of locomotor speed. The direction heading between these two points was used to identify rotations and change in direction. Rotations were quantified as a continuous 180°turn containing no more than 90 degrees of continuous rotation in the opposite direction.

### Brain slice patch clamp electrophysiology and optogenetic stimulation experiments

Enhancer AAV injected mice were deeply anaesthetized by intraperitoneal administration of Avertin (20 mg kg ^1^) and were perfused through the heart with carbogenated (95% O2/5% CO2) artificial aCSF consisting of (in mM): 92 N-methyl-D-glucamine (NMDG), 2.5 KCl, 1.25 NaH2PO4, 30 NaHCO3, 20 4-(2-hydroxyethyl)-1-piperazineethanesulfonic acid (HEPES), 25 glucose, 2 thiourea, 5 Na-ascorbate, 3 Na-pyruvate, 0.5 CaCl2·4H2O and 10 MgSO4·7H2O. (VT1200S, Leica Biosystems or Compresstome VF-300, Precisionary Instruments) using a zirconium ceramic blade and following the NMDG protective recovery method [Ting 2014]. Mouse brains were sectioned in the coronal plane such that the angle of slicing was perpendicular to the pial surface. After sections were obtained, slices were transferred to a warmed (32-34 °C) initial recovery chamber filled with NMDG aCSF under constant carbogenation. After 12 min, slices were transferred to a chamber containing HEPES holding aCSF solution consisting of (in mM): 92 NaCl, 2.5 KCl, 1.25 NaH2PO4, 30 NaHCO3, 20 HEPES, 25 glucose, 2 thiourea, 5 sodium ascorbate, 3 sodium pyruvate, 2 CaCl2·4H2O and 2 MgSO4·7H2O, continuously bubbled with 95% O2/5% CO2. Slices were held in this chamber until use in acute patch clamp recordings or until fixed in 4% PFA for later histological processing.

Brain slices were placed in a submerged, heated (32-34°C) chamber that was continuously perfused with fresh, carbogenated aCSF consisting of (in mM): 119 NaCl, 2.5 KCl, 1.25 NaH2PO4, 24 NaHCO3, 12.5 glucose, 2 CaCl2·4H2O and 2 MgSO4·7H2O (pH 7.3-7.4). Neurons were visualized with an upright microscope (Scientifica) equipped with infrared differential interference contrast (IR-DIC) optics and both 4x air and 40x water immersion objectives as well as epifluorescence illumination and filter cubes to detect SYFP2, mTFP1, and tdTomato fluorescence. Glass patch clamp pipettes were pulled to an open tip resistance of 2-6 MW when filled with the internal recording solution consisting of (in mM): 110.0 K-gluconate, 10.0 HEPES, 0.2 EGTA, 4 KCl, 0.3 Na2-GTP, 10 phosphocreatine disodium salt hydrate, 1 Mg-ATP, 20 mg/mL glycogen, 0.5U/mL RNase inhibitor (Takara, 2313A), 0.5% biocytin and 0.02 Alexa 594 or 488 pH adjusted to 7.3 with KOH. Whole cell somatic recordings were acquired using a Multiclamp 700B amplifier and custom acquisition software written in Igor Pro (MIES https://github.com/AllenInstitute/MIES. Electrical signals were digitized at 50 kHz by an ITC-18 (HEKA) and were filtered at 10 kHz. The pipette capacitance was compensated, and the bridge was balanced during the current clamp recordings.

In triple RO injection experiments, recorded striatal interneurons in the dorsal striatum region were stimulated by injecting a sequence of 1s hyperpolarizing and depolarizing square wave current steps varying from −200pA to 500pA (D1 MSN and cholinergic), −100pA to 225pA (Sst-Chodl) and −300pA to 675pA (Pvalb-Pthlh)in 50pA, 25pA and 75pA increments for cholinergic Sst-Chodl and Pvalb-Pthlh respectively. Intrinsic electrophysiological features were extracted using Python-based code adapted from Intrinsic Physiology Feature Extractor (IPFX, https://github.com/AllenInstitute/ipfx).

Electrophysiological features used to cluster cells in 3D plots and highlight their physiological differences were calculated as follows: Frequency/Current (FI) Slope: calculated as the initial linear slope of the firing rate as a function of the current injection amplitude; Steady-state input resistance: calculated from the linear fit of the current-voltage relationship measured in response to the series of hyperpolarizing steps; Upstroke/downstroke ratio: ratio of the maximum dVdt to the minimum dVdt for the first spike during the first current step that evoked spiking; Resting membrane potential: calculated as the membrane potential voltage with no current bias or current pulse injected; Action potential(AP) half-width: calculated as the duration of the action potential at the voltage halfway between AP threshold and the AP peak.

For STX injection experiments measuring peak photocurrent amplitude of ChR2-expressing D1 MSNs for various expression strategies, the area with maximal YFP reporter native fluorescence was first identified. As labeling appeared as dense YFP+ neuropil rather than clearly discernable individual MSN somata, neurons within this area with healthy appearing MSN-like morphology under IR-DIC were selected for voltage clamp recordings with a holding potential set to −70mV. Blue light was delivered from a mercury arc lamp attached to light guide directed through the 40X microscope objective equipped with blue excitation spectrum bandpass filter cube. The power density was empirically set to illicit maximal or saturating photocurrent response amplitude. Recorded neurons with no measurable peak photocurrent in response to blue light stimulation were presumed as uninfected or putative D2 MSNs and were excluded from the analysis. Putative D1 MSNs exhibited clear photocurrent responses to 1s blue light stimulation (minimal 3 stimulation sweeps recorded per cell), and the peak photocurrent was measured at the onset of the response (within a 5ms window). The opsin variant ChR2(CRC)-EYFP was selected to match to in vivo optogenetics experiments in this study, and notably this variant shows step-like activation kinetics with minimal to no desensitization relative to strong desensitization observed with other variants including ChR2(H134R)-EYFP. Recordings in the current clamp mode were obtained for select neurons to measure FI curves to support MSN identity and signature firing properties, as well as for measuring blue light evoked action potential firing from the rest.

### Macaque *in vivo* enhancer AAV injections and histology

All procedures used with macaque monkeys conformed to the guidelines provided by the US National Institutes of Health and were approved by the University of Washington Animal Care and Use Committee. Two southern pig-tailed macaques (Macaca nemestrina) were used in this study for single and duplexed *in vivo* injection of iodixanol gradient purified enhancer AAV vectors using the Brainsight® robotic surgery system (Rogue Research) (animal one, a 7-year-old 5.8 kg female, and animal two, a 6-year-old 15kg male, respectively). Brain MRIs for each animal were acquired and utilized for pre-planning of virus injection surgery using the Brainsight**^®^** vet robot software v2.5 (Rogue Research). On the day of surgery, the animals were deeply anesthetized and positioned with the head in the stereotaxic frame of the Brainsight**^®^** surgery platform. A skin incision was made to expose the top of the skull, and burr hole craniotomies were drilled over the planned injection trajectory locations under the guidance of the Brainsight**^®^** robotic surgery arm. After craniotomy, a prefabricated steel cannula with PE tubing attachment^100^ was loaded with a total of 20 L AAV vector solution and attached to the robotic arm. Animal one received two distinct single AAV treatments: 1) In the right hemisphere, a total of 5uL of virus was injected into caudate, and 5uL into putamen using two distinct injection tracks with AAV vector for enhancer AiE0780m driving SYFP2 (AiP12610). 2) In the left hemisphere, a total of 5uL of virus was injected into putamen with AAV vector for enhancer AiE0452h driving mTFP1 (AiP12700). Animal two received one duplexed AAV treatment. A total 5uL of virus was injected into putamen using a mix of the two AAV vectors. The diluted titer for these injections was 2.00E+13 vg/mL. For each injection track, the cannula was advanced to the starting point of the injection target. Then, 1uL of virus was gradually injected at each of five depths with 1mm spacing as the cannula tip was retracted dorsally. At the end of the final injection point in each track, the cannula was left in place for 10 min to allow virus diffusion prior to extracting the cannula from the brain. Note that additional virus injection sites were made in other distal brain locations in each macaque, but striatal labeling remained physically clearly separated from distal injection sites.

At 35-40 days post-injection, each animal was sacrificed for brain collection and tissue expression analysis. The animals were transcardially perfused with 2L NMDG-aCSF solution. The brain was then removed and rapidly transported from the WaNPRC to the Allen Institute in chilled NMDG-aCSF solution for further tissue processing. The brain was first hemisected and drop-fixed in freshly prepared 4% PFA in PBS for 48 hours at 4°C. After PFA fixation, the brain was transferred into 1XPBS + 0.01% sodium azide solution. The brain hemispheres were examined and then 0.5 cm thick coronal slabs were cut through the striatum regions. Serial brain slabs were trimmed to fit into the wells of a six well tissue culture plate and submerged in 30% sucrose solution for a minimum of 48hrs before further processing for histological analysis. Tissue slabs were mounted on the stage of a freezing-sliding microtome (Leica model SM2010R) on a bed of OCT and sub-sectioned to 30m thickness and stored in 1XPBS + 0.01% sodium azide solution. Sections in the region of interest were selected for propidium iodide and DAPI staining and then slide mounted on 2×3 glass slides. Alternately, some tissue sections were used for free-floating immunostaining using the following antibodies: 1) chicken anti-GFP IgY (1:2000), mouse anti-DARPP32 IgG1 (1:500-2000), and rat anti-mTFP IgG (1:1000). Secondary antibodies were as follows: Alexa-488 conjugated goat anti-chicken IgY (1:1000), Alexa-647 conjugated goat anti-mouse IgG1 (1:1000), and BV480 conjugated goat anti-rat IgG (1:1000). Sections were dried onto slides on a slide warmer and cover-slipped with Vectashield Hardset mounting medium (Vector Labs) for PI and DAPI plus native SYFP2 imaging or Prolong Gold Antifade mounting medium (Life Technologies) for antibody staining experiments. Plea

## RESOURCE AVAILABILITY

Enhancer characterization data is available to view at the Allen Institute for Brain Science Genetic Tools Atlas (GTA): https://portal.brain-map.org/genetic-tools/genetic-tools-atlas

Mouse brain expression data from this study can be accessed in the GTA using the publication tag Hunker, Wirthlin et al **Table S2** provides a manifest of experiments from this study in GTA and can serve as a guide to searching for specific datasets directly (e.g., by Donor ID, Vector ID, Enhancer ID, etc.). Additional data types will be added with future data releases to the GTA.

SSv4 scRNA-seq data are available at the Neuroscience Multi-omic Archive (NeMO): https://data.nemoarchive.org/other/grant/uf1_tasic/tasic/transcriptome/scell/SSv4/mouse/raw/

Primary screen epifluorescence and Serial-two-photon tomography (STPT) data are available at Brain Image Library (BIL) at: https://doi.brainimagelibrary.org/doi/10.35077/g.1159

Primary screen epifluorescence data can be found at BIL at: https://download.brainimagelibrary.org/77/43/774317f96671fbd7/

Serial-two-photon tomography (STPT) data can be found at BIL at: https://download.brainimagelibrary.org/07/7d/077dbfded4d55b5e/

Plasmid DNA for enhancer AAV vectors have been deposited to Addgene (see Key Resources table) as part of the NIH BRAIN Armamentarium collection (https://www.addgene.org/collections/brain-armamentarium/) and are available for distribution under standard UBMTA. Users can filter for a list of viral aliquot offerings from the Addgene Armamentarium page at the link above by entering Ting table field. Plasmid Vector IDs and Enhancer IDs used in each main and supplementary Figure are summarized in **Table S3**. Further information, including mFISH data and requests for resources and reagents should be directed to and will be fulfilled by the lead contact, Jonathan Ting.

## ACKNOWLEDGMENTS

We wish to thank the Allen Institute founder, Paul G. Allen, for his vision and the NIH BRAIN Initiative Armamentarium for Precision Brain Cell Access (https://braininitiative.nih.gov/armamentarium). This research was supported by U.S. National Institutes of Health (NIH) BRAIN Initiative Armamentarium Grant UF1MH128339 (BTa, JTT, BPL, TLD, and TEB) and in part by NIH BRAIN Initiative Human and Mammalian Brain Atlas (HMBA) BICAN grant UM1MH130981 (ESL, HZ), NIH grant R01MH123620 (BEK), and Allen Institute funding to the Genetic Tools project. We thank additional members of the core facilities and joint Brian Science shared resource teams including Laboratory Animal Services, Transgenic Colony Management Team, Animal Care Team, Neurosurgery and Behavior Team, Viral Technology Team, Data and Technology Team, Histology Team, Imaging Team, Tissue Processing Team, Project Management Team, and Facilities Team. We thank the WaNPRC surgery and veterinary support staff and Chris English for necropsy support. The WaNPRC is supported by the NIH Office of Research Infrastructure Programs (ORIP) under award number P51OD010425 and U420D011123. We thank Dr. Elizabeth Buffalo for access to the BrainSight injection robot system and Megan Jutras for BrainSight support and training. We thank DISC lab and Tim Wilbur at the University of Washington for support with macaque MRI imaging. We thank TL Wong from the Kojima lab for support with fabricating virus injection cannulae and Shane Gibson from the Horwitz lab for support with macaque surgery and sample processing. Illustrations in Figure 6 were created with BioRender.com.

## AUTHOR CONTRIBUTIONS

<u>Conceptualization:</u> ACH, MEW, SFO, BPL, TLD, BTa, TEB, JTT. <u>Methodology:</u> ACH, MEW, GG, NJJ, MH, VO, SV, NL, NT, NW, WDL, YMB, JLB, BBG, XOA, RAM, SH, JKM, MJT, AW, ND, JeA, YK, GH, SFO, BPL, TLD, BTa, TEB, JTT. <u>Software:</u> MH, JRA, AdA, JoA, CB, PB, ABC, RC, KC, SFD, PD, TF, JG, ZH, NAL, ZM, JeM, JM, EM, ALO, CRe, DBR, LSa, SS, YW, LLN, BK, FC. <u>Validation:</u> ACH, VO, NT, NW, SWW, BTh, JKM, SFO, BPL, TLD, BTa, JTT. <u>Formal Analysis:</u> ACH, MEW, GG, NJJ, VO, SV, NL, NT, WDL, YBS, EM, SFO, JTT. <u>Investigation:</u> ACH, GG, VO, SV, NL, NT, NW, WDL, YMB, JLB, BBG, AvA, APA, SB, DB, KB, NID, TCE, AG, CG, WH, TJ, ZJ, JK, HL, JoM, EM, RN, KiN, KaN, BO, AAO, THP, CAP, LP, SR, CRi, ARS, AR, MJT, MT, JWi, JeA, YK, GH, SFO, JTT. <u>Resources:</u> MEW, MH, XOA, RAM, TB, JB, GB, BC, TC, MC, TD, MD, NPD, ELG, OH, WVH, CH, DLJ, ML, EL, RM, DN, ALO, NP, EP, MR, DFR, KR, JS, NVS, LSh, TW, TZ, LE, SY, FC, SM, KAS, JWa, IE, MP, JTT. <u>Data</u> <u>curation:</u> ACH, MEW, GG, NJJ, MH, SV, NL, WDL, YBS, SWW, BTh, PMB, PB, DD, NAL, EM, IE, MP, SFO, BPL, TLD, JTT. <u>Writing - original draft:</u> ACH, MEW, GG, NJJ, SFO, JTT. <u>Writing - review and</u> <u>editing:</u> ACH, MEW, MJT, BK, SFO, BPL, TLD, BTa, TEB, JTT. <u>Visualization:</u> ACH, MEW, GG, NJJ, MH, SV, JLB, SO, REAS, SFD, SL, NAL, SMo, CRe, NS, BS, LLN, FC, SFO, JTT. <u>Supervision:</u> BBG, YBS, SH, MJT, AW, ND, LLN, SY, JeA, FC, SM, KAS, JWa, HZ, ESL, YK, GH, SFO, BPL, TLD, BTa, TEB, JTT. <u>Project Administration:</u> SWW, BTh, SO, REAS, KG, TM, LE, IE, MP, GH, BPL, TLD, BTa, TEB, JTT. <u>Funding acquisition:</u> BK, HZ, ESL, BPL, TLD, BTa, TEB, JTT.

## SUPPLEMENTAL INFORMATION

**Document S1. Supplemental Figures S1 S10**

**Table S1. Detailed enhancer metadata**

**Table S2. GTA experiment metadata**

**Table S3. List of AAV vector plasmid IDs and enhancer IDs used in each figure**

