## Supplementary material for "Enhancer AAV toolbox for accessing and perturbing striatal cell types and circuits": Table_S3

**Table S3:** List of AAV vector plasmid IDs and enhancer IDs used in each figure

| **Figure #** | **Plasmid_ID (Enhancer_ID)** |
| --- | --- |
| Figure 1 | AiP12237 (AiE0452h) |
| Figure 2 | AiP12237 (AiE0452h), AiP12410 (AiE0603m), AiP12609 (AiE0779m), AiP12689 (AiE0682h), AiP12787 (AiE0140h_3xC2), AiP13038 (AiE0743m_3xC2) |
| Figure 3 | AiP12229 (AiE0441h), AiP12231 (AiE0445h), AiP12233 (AiE0447h), AiP12236 (AiE0450h), AiP12237 (AiE0452h), AiP12408 (AiE0600m), AiP12410 (AiE0603m), AiP12446 (AiE0616h), AiP12447 (AiE0617h), AiP12451 (AiE0621h), AiP12514 (AiE0367h_C2), AiP12555 (AiE0367h_3xC2), AiP12602 (AiE0769m), AiP12609 (AiE0779m), AiP12610 (AiE0780m), AiP12631 (AiE0743m), AiP12689 (AiE0682h), AiP12701 (hChATp), AiP12747 (AiE0784m), AiP12787 (AiE0140h_3xC2), AiP12982 (AiE0452h_3xC2), AiP13038 (AiE0743m_3xC2), AiP13044 (AiE0779m_3xC2), AiP13445 (AiE0873m), AiP13792 (AiE0951h), AiP13905 (AiE0441h_3xC2), AiP14428 (AiE1191m), AiP14496 (AiE0873m_3xC2), AiP14696 (AiE0779m_6xC2), AiP15028 (AiE1404m), AiP15050 (AiE1426m), AiP15043 (AiE1419m), AiP15045 (AiE1421m), AiP1530 (AiE2116m_3xC2) |
| Figure 4 | AiP12237 (AiE0452h), AiP12602 (AiE0769m), AiP12609 (AiE0779m), AiP12610 (AiE0780m), AiP12689 (AiE0682h), AiP12787 (AiE0140h_3xC2), AiP13038 (AiE0743m_3xC2), AiP13278 (AiE0779m_3xC2), AiP14035 (AiE0452h_3xC2), AiP14134 (AiE0452h_3xC2), AiP14496 (AiE0873m_3xC2), AiP14496 (AiE0873m_3xC2), AiP15050 (AiE1426m), AiP5043 (AiE1419m) |
| Figure 5 | AiP13781 (AiE0452h), AiP14825 (AiE0441h_3xC2), AiP15260 (AiE0779m_3xC2), AiP15578 (AiE0780m) |
| Figure 6 | AiP13278 (AiE0779m_3xC2), AiP13044 (AiE0779m_3xC2) |
| Figure 7 | AiP12609 (AiE0779m), AiP12689 (AiE0682h), AiP12700 (AiE0452h), AiP13738 (AiE0743m_3xC2), AiP13808 (AiE0140h_3xC2), AiP15050 (AiE1426m) |
| Figure 8 | AiP12229 (AiE0441h), AiP12233 (AiE0447h), AiP12237 (AiE0452h), AiP12514 (AiE0367h_C2), AiP12609 (AiE0779m), AiP12610 (AiE0780m), AiP13905 (AiE0441h_3xC2) |
| Figure S1 | Bioinformatic analysis on enhancers only: (AiE1426m), (AiE1191m), (AiE0784m), (AiE1190m), (AiE0785m), (AiE1419m), (AiE0869m), (AiE0603m), (AiE1404m), (AiE0743m), (AiE0600m), (AiE0873m), (AiE0779m), (AiE0782m), (AiE0780m), (AiE1421m), (AiE0868m), (AiE0951h), (AiE0617h), (AiE0450h), (AiE0621h), (AiE0444h), (AiE0612h), (AiE0367h), (AiE0441h), (AiE0452h), (AiE0445h), (AiE0618h), (AiE0682h), (AiE0351h), (AiE0616h), (AiE0447h), (AiE1026h) |
| Figure S2 | AiP12421 (AiE0444h), AiP12448 (AiE0618h), AiP1530 (AiE2116m_3xC2), AiP12446 (AiE0616h), AiP12447 (AiE0617h), AiP13868 (AiE1026h) |
| Figure S3 | AiP12013 (AiE0351h), AiP13566 (AiE0351h_3xC2), AiP12025 (AiE0367h), AiP12514 (AiE0367h_C2), AiP12555 (AiE0367h_3xC2), AiP12229 (AiE0441h), AiP13905 (AiE0441h_3xC2), AiP12233 (AiE0447h), AiP12236 (AiE0450h), AiP12410 (AiE0603m), AiP12446 (AiE0616h_3xC2), (AiE0618h), AiP12451 (AiE0621h), AiP13344 (AiE0621h_3xC2), AiP12626 (AiE0782m), AiP12747 (AiE0784m), AiP12614 (AiE0785m), AiP13868 (AiE1026h), AiP14428 (AiE1191m), AiP12446 (AiE0616h), AiP12609 (AiE0779m), AiP13044 (AiE0779m_3xC2), AiP14696 (AiE0779m_6xC2), AiP12610 (AiE0780m), AiP12421 (AiE0444h), AiP12231 (AiE0445h), AiP12237 (AiE0452h), AiP12982 (AiE0452h_3xC2), AiP12442 (AiE0612h), AiP12447 (AiE0617h), AiP14427 (AiE1190m), AiP13048 (AiE0576h_3xC3), AiP12631 (AiE0743m), AiP13038 (AiE0743m_3xC2), AiP13362 (AiE0868m), AiP13363 (AiE0869m), AiP13445 (AiE0873m), AiP14496 (AiE0873m_3xC2), AiP12408 (AiE0600m), AiP12689 (AiE0682h), AiP12602 (AiE0769m), AiP15028 (AiE1404m), AiP15050 (AiE1426m), AiP12787 (AiE0140h_3xC2), AiP13792 (AiE0951h), AiP15043 (AiE1419m), AiP15045 (AiE1421m) |
| Figure S4 | AiP14452 (AiE0351h), AiP14448 (AiE0367h), AiP14449 (AiE0367h_3xC2), AiP14453 (AiE0441h), AiP14458 (AiE0441h_3xC2), AiP14451 (AiE0447h), AiP14450 (AiE0450h), AiP14455 (AiE0526h), AiP14456 (AiE0603m), AiP14454 (AiE783m), AiP14457 (AiE0784m), AiP14459 (AiE0785m), AiP14460 (AiE0452h_3xC2), AiP14461 (AiE0779m_3xC2) |
| Figure S5 | AiP14696 (AiE0779m_6xC2), AiP13044 (AiE0779m_3xC2), AiP12609 (AiE0779m), AiP12610 (AiE0780m), AiP12421 (AiE0444h), AiP12231 (AiE0445h), AiP12982 (AiE0452h_3xC2), AiP12237 (AiE0452h), AiP2555 (AiE0367h_3xC2), AiP12514 (AiE0367h_C2), AiP13905 (AiE0441h_3xC2), AiP12229 (AiE0441h), AiP12233 (AiE0447h), AiP12236 (AiE0450h), AiP12410 (AiE0603m), AiP12747 (AiE0784m), AiP14428 (AiE1191m), AiP13038 (AiE0743m_3xC2), AiP12631 (AiE0743m), AiP14496 (AiE0873m_3xC2), AiP13445 (AiE0873m), AiP12701 (hChATp) |
| Figure S6 | AiP12229 (AiE0441h), AiP13905 (AiE0441h_3xC2), AiP12451 (AiE0621h), AiP12609 (AiE0779m), AiP13044 (AiE0779m_3xC2), AiP14696 (AiE0779m_6xC2), AiP12446 (AiE0616h), AiP12610 (AiE0780m), AiP12237 (AiE0452h), AiP12982 (AiE0452h_3xC2), AiP12421 (AiE0444h), AiP12447 (AiE0617h), AiP12787 (AIE0140h_3xC2), AiP13792 (AiE0951h), AiP15043 (AiE1419m,) AiP15045 (AiE1421m), AiP12408 (AiE0600m), AiP12689 (AiE0682h), AiP12602 (AIE0769m), AiP15050 (AiE1426m), AiP12631 (AiE0743m), AiP13038 (AiE0743m_3xC2), AiP13445 (AiE0873m), AiP14496 (AiE0873m_3xC2), AiP12701 (hChATp) |
| Figure S7 | iCre vectors: AiP13777 (AiE0682h), AiP1793 (AiE0600m), AiP13780 (AiE0743m_3xC2), AiP15508 (AiE1419m), AiP13778 (AiE0450h), AiP13781 (AiE0452h), AiP13779 (AiE0779m_3xC2), AiP15578 (AiE0780m); FlpO vectors: AiP14826 (AiE0441h_3xC2), AiP15100 (AiE0140h_3xC2), AiP14201 (AiE0779m_3xC2), AiP14573 (AiE0452h) |
| Figure S8 | AiP13278 (AiE0779m_3xC2), AiP15578 (AiE0780m),  AiP11763 (pAAV-hEF1a-DIO-ChR2(CRC)-EYFP-WPRE-HGHpA) |
| Figure S9 | AiP12233 (AiE0447h), AiP12542 (AiE0410m), AiP13738 (AiE0743m_3xC2), AiP13808 (AiE0140h_3xC2), AiP15050 (AiE1426m) |
| Figure S10 | AiP12700 (AiE0452h), AiP12610 (AiE0780m) |
