## Supplementary material for "Enhancer AAV toolbox for accessing and perturbing striatal cell types and circuits": Document_S1_Supplementary_Figures

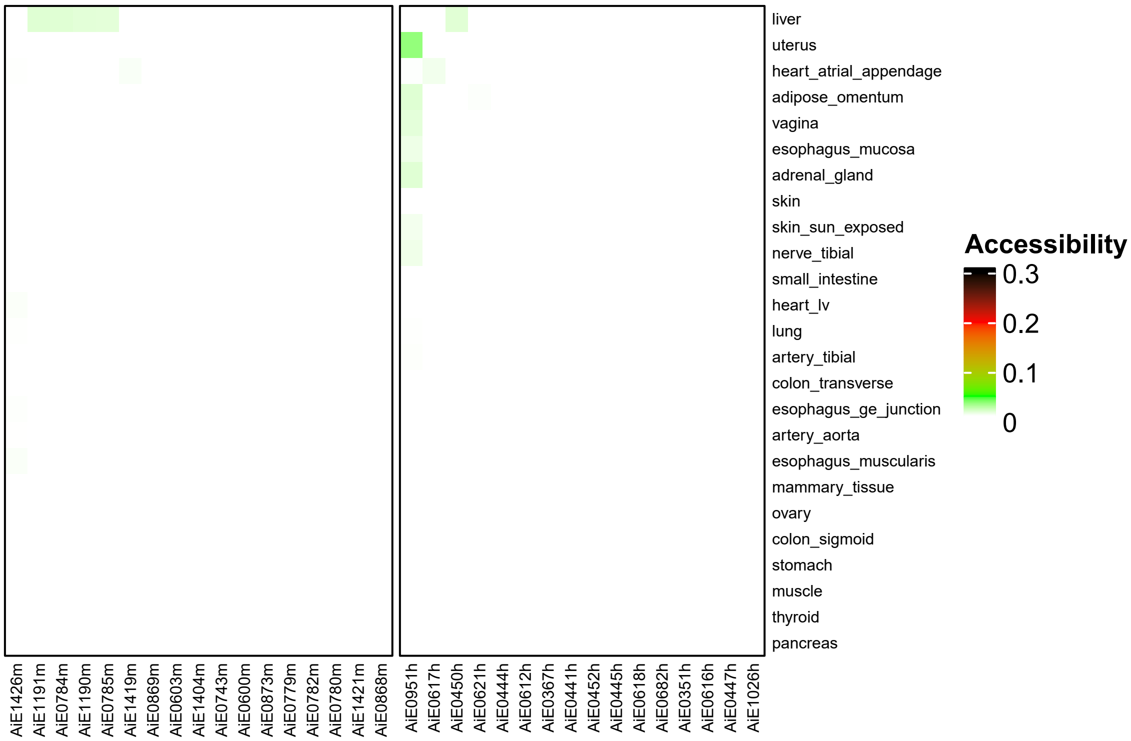


**Figure S1**: Predictability of enhancer AAV expression across human body organs.
Single-cell open chromatin profiles were grouped within each tissue into pseudo-bulk aggregates, then normalized according to the signal (reads in peaks) within the dataset. Accessibility scores for all enhancers found in ATAC-seq datasets (except for AiE0769m which did not have hg38 liftover coordinates) were computed using multiBigwigSummary command from DeepTools. All tested enhancers had low accessibility scores predicting little to no enhancer activities within each non-brain body tissue listed.

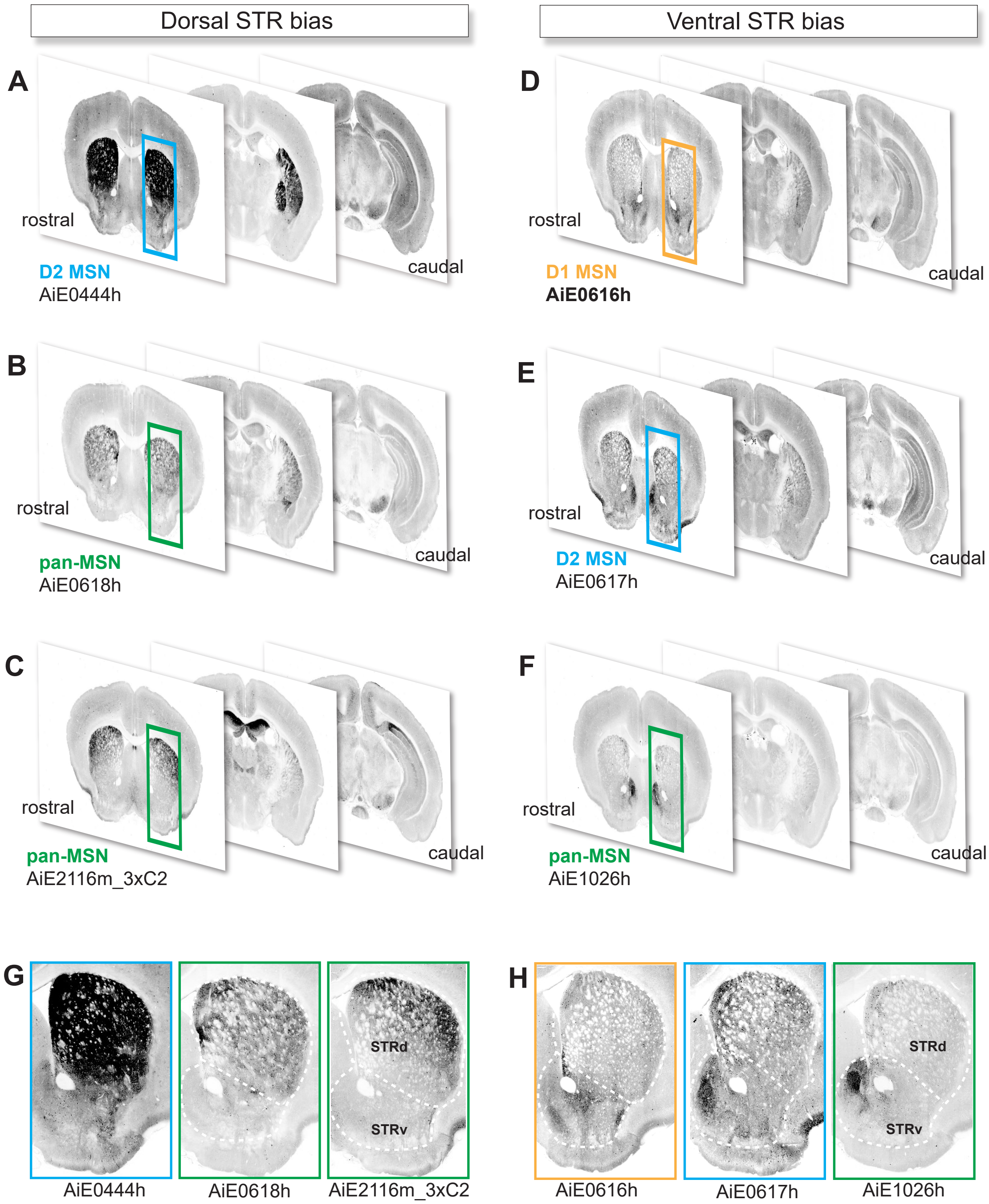


**Figure S2:** Enhancer AAVs with dorsal or ventral striatal expression. A-C) Selected coronal STPT images of enhancer AAVs with dorsal striatum bias. A) Illustrates D2 MSN enhancer AiE0444h, B-C) Illustrates MSN enhancers AiE0618h and AiE2116m_3xC2. D-F) Selected coronal STPT images of enhancer AAVs with ventral striatum bias. D) D1 MSN enhancer AiE0616h and E) D2 MSN enhancer AiE0617h and F) MSN enhancer AiE1026h. G-H) Zoomed in views showing side-by-side expression patterns comparing G) dorsal-biased versus H) ventral-biased enhancer AAVs..

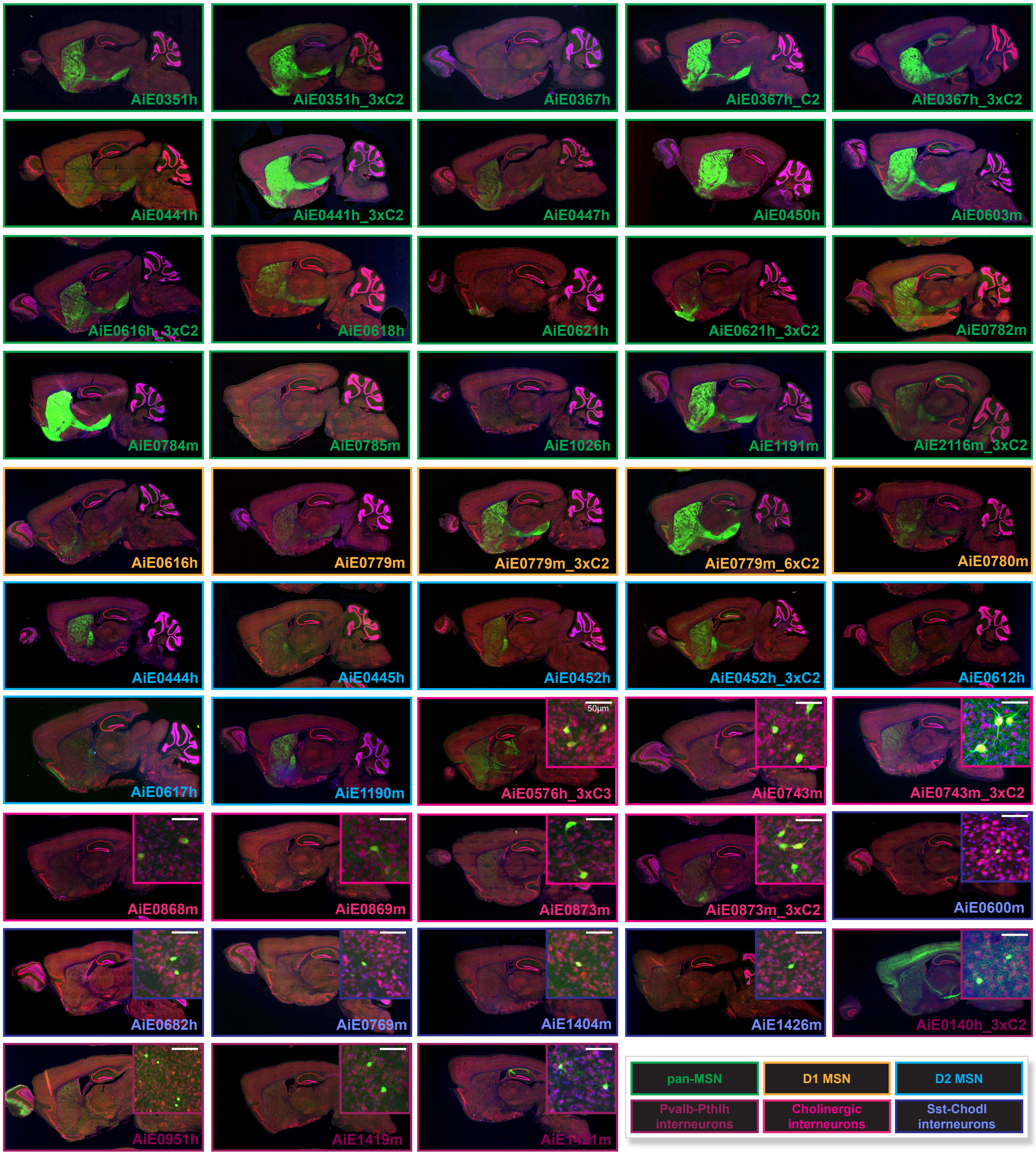


**Figure S3**: Primary screening results for all best-in-class striatal enhancer AAVs. Sagittal brain sections with propidium iodine (red), DAPI (blue), and SYFP2 (green) native fluorescence for each enhancer driving SYFP2 included in this study. Colored boxes around sagittal sections correspond to cell type specificity of each enhancer AAV. Images were not acquired under matched conditions and have been adjusted to optimally highlight brain wide expression patterns for enhancer specificity comparison purposes.

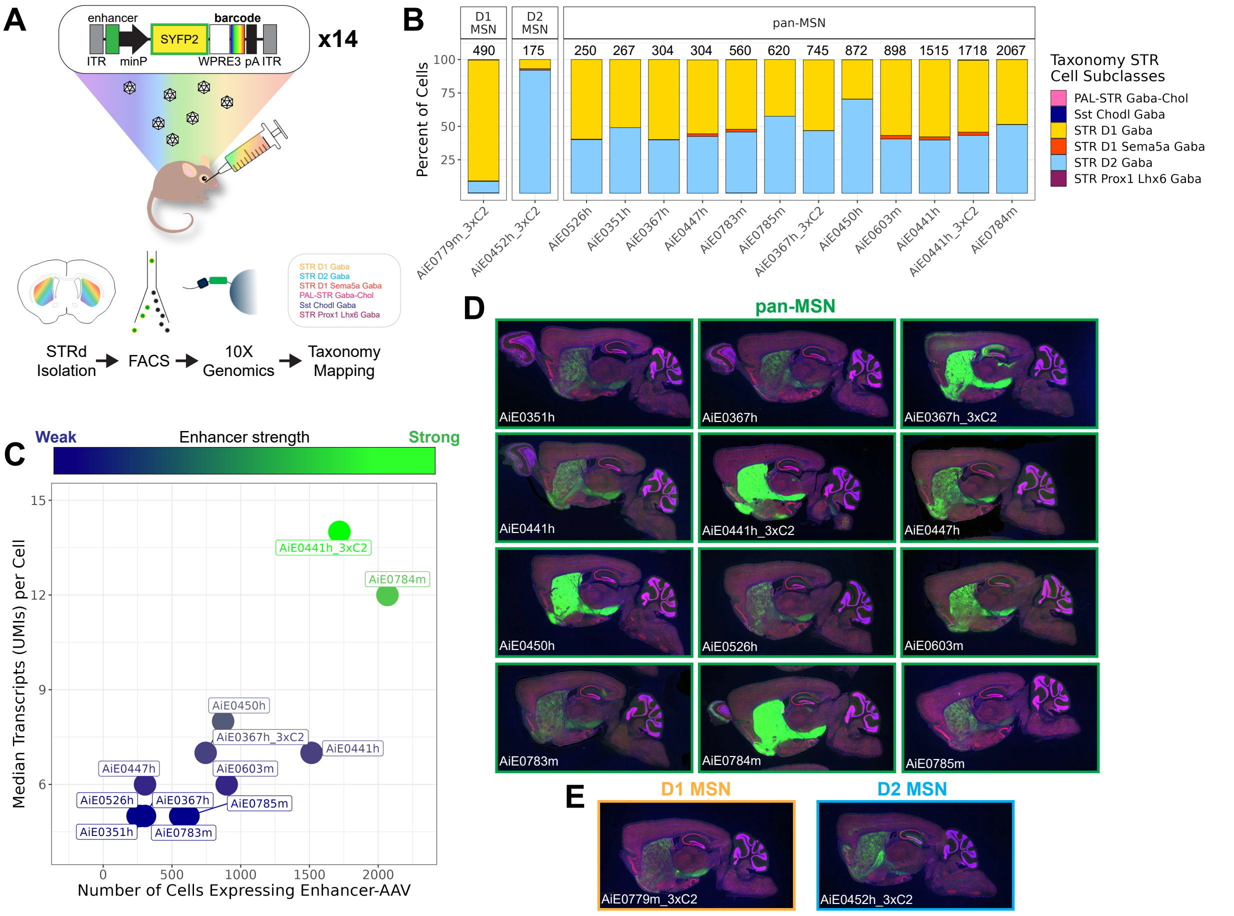


**Figure S4**: Comparison of pan-MSN enhancer AAVs using barcoded multiplex scRNA-seq.

A) Diagram of the experimental design. Twelve pan-MSN enhancers (plus one D1 MSN enhancer and one D2 MSN enhancer as controls) were paired with a unique 8bp barcode, package separately into AAVs and injected RO as a pool into a single adult mouse. SYFP2+ cells were sorted and prepared for scRNA-seq using 10X Genomics v3.1. Cells were mapped onto the 10x Mouse Whole Brain taxonomy (CCN20230722) to determine cell subclasses that express each enhancer. B) Proportion of cells that label each of the targeted striatal cell subclasses. The labels above the columns represent the total number of cells expressing each individual enhancer. C) Comparison of number of cells expressing each pan-MSN enhancer (measure of completeness of labeling) with the median number of enhancer-driven transcripts per cell (measure of enhancer strength). Kruskal-Wallis rank sum test was used to assess differences in UMIs by enhancer (chi-squared = 1163.3, df = 11, $p < 2.2e-16$). Dunn's test revealed significant differences between many of the enhancers. Two enhancers AiE0441h_3xC2 and AiE0784m showed significantly stronger expression than all other enhancers ($p<0.05$) but were not significantly different from each other. D-E) Epifluorescence images of each of the barcoded enhancer AAV vectors injected RO individually into adult mice. Each section was stained with Propidium iodine (red) and DAPI (blue). SYFP2 (green) is native fluorescence.

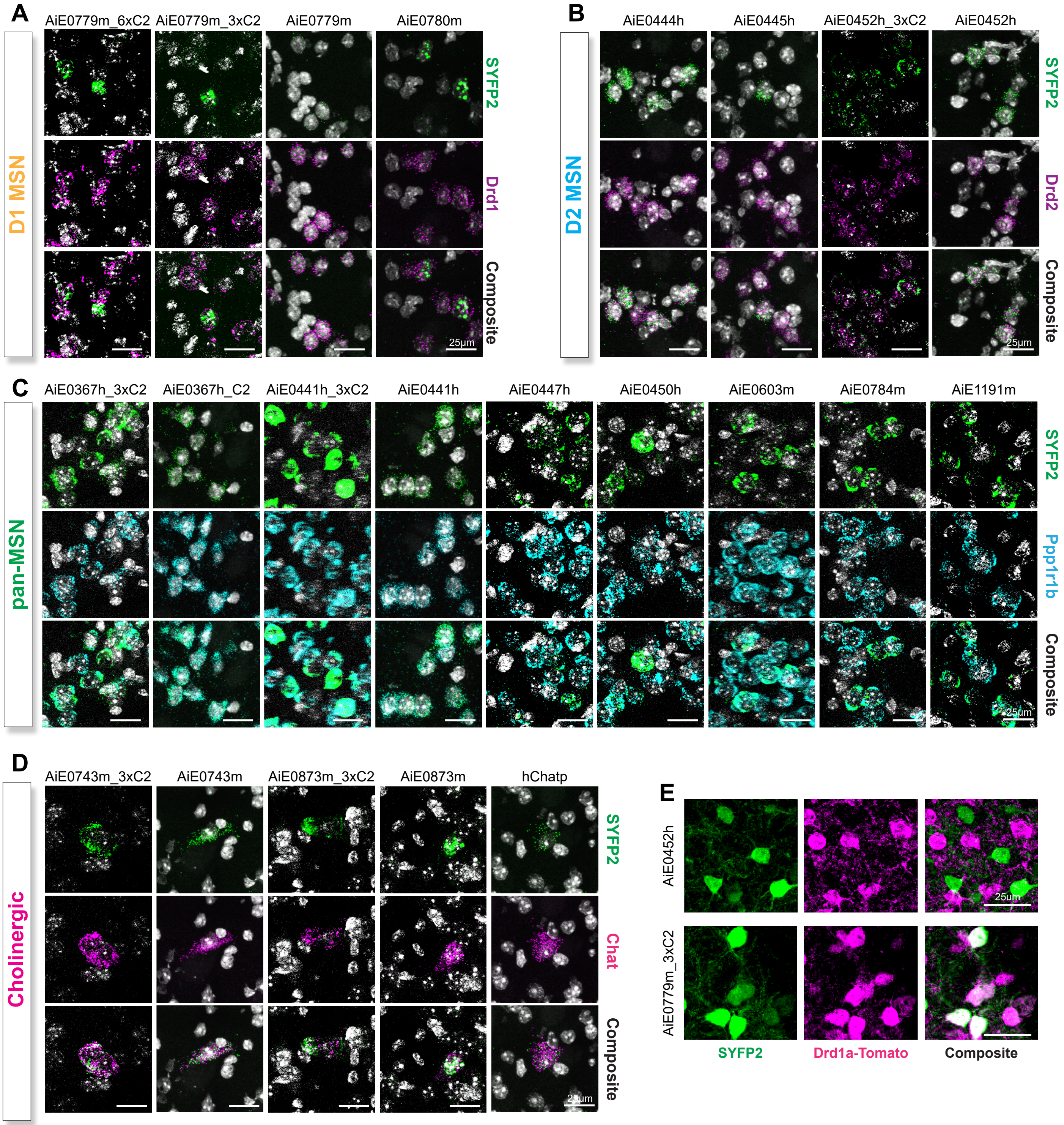


**Figure S5**: RNAscope and IHC results for quantifying enhancer on-target specificities.
A) D1 MSN enhancers with probes against SYFP2 mRNA (top row) and Drd1 (middle row). B) D2 MSN enhancers with probes against SYFP2 mRNA (top row) and Drd2 (middle row). C) Pan-MSN enhancers with probes against SYFP2 mRNA (top row) and Ppp1r1b (middle row). D) Cholinergic enhancers with probes against SYFP2 mRNA (top row) and Chat (middle row). E) IHC results for quantifying D1 MSN enhancer AiE0779m_3xC2 and D2 MSN enhancer AiE0452h using SYFP2 native fluorescence (left) and tdTomato fluorescence (middle) following RO injection of AAV vectors into Drd1a-tdTomato hemizygous mice.

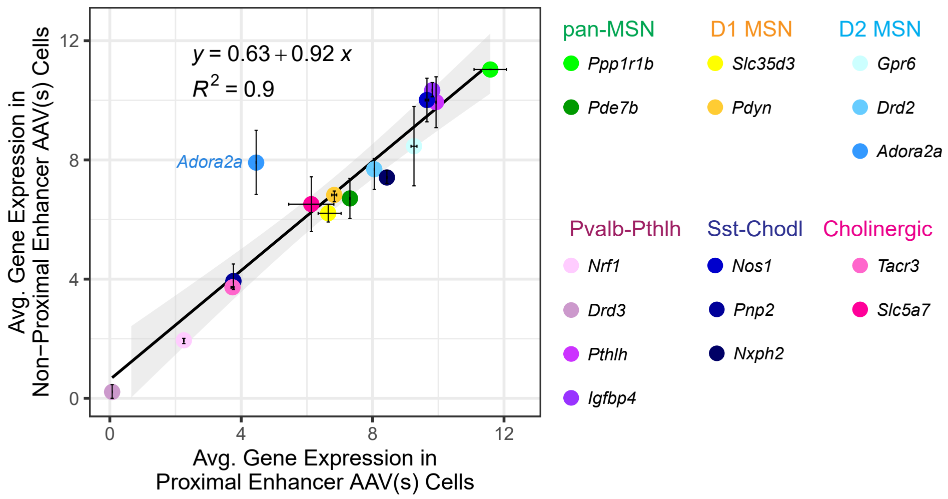


**Figure S6**: Enhancer AAVs do not cause down regulation of endogenous marker gene transcripts. Comparison of normalized average gene expression (log2(CPM+1)) from sc-RNAseq datasets (SSv4) for select marker genes found proximal (x-axis) or non-proximal (y-axis) for each cell type target. Error bars represent min and max average gene expression across proximal or non-proximal enhancer AAVs targeting the same cell type. R2=0.9 indicates strong correlation between average marker gene expression in proximal vs non-proximal enhancers. However, the Studentized residual analysis identified Adora2a as a potential outlier, with a residual value of 7, indicating its expression deviates significantly from the predicted regression trend. Please see Methods section for more details.

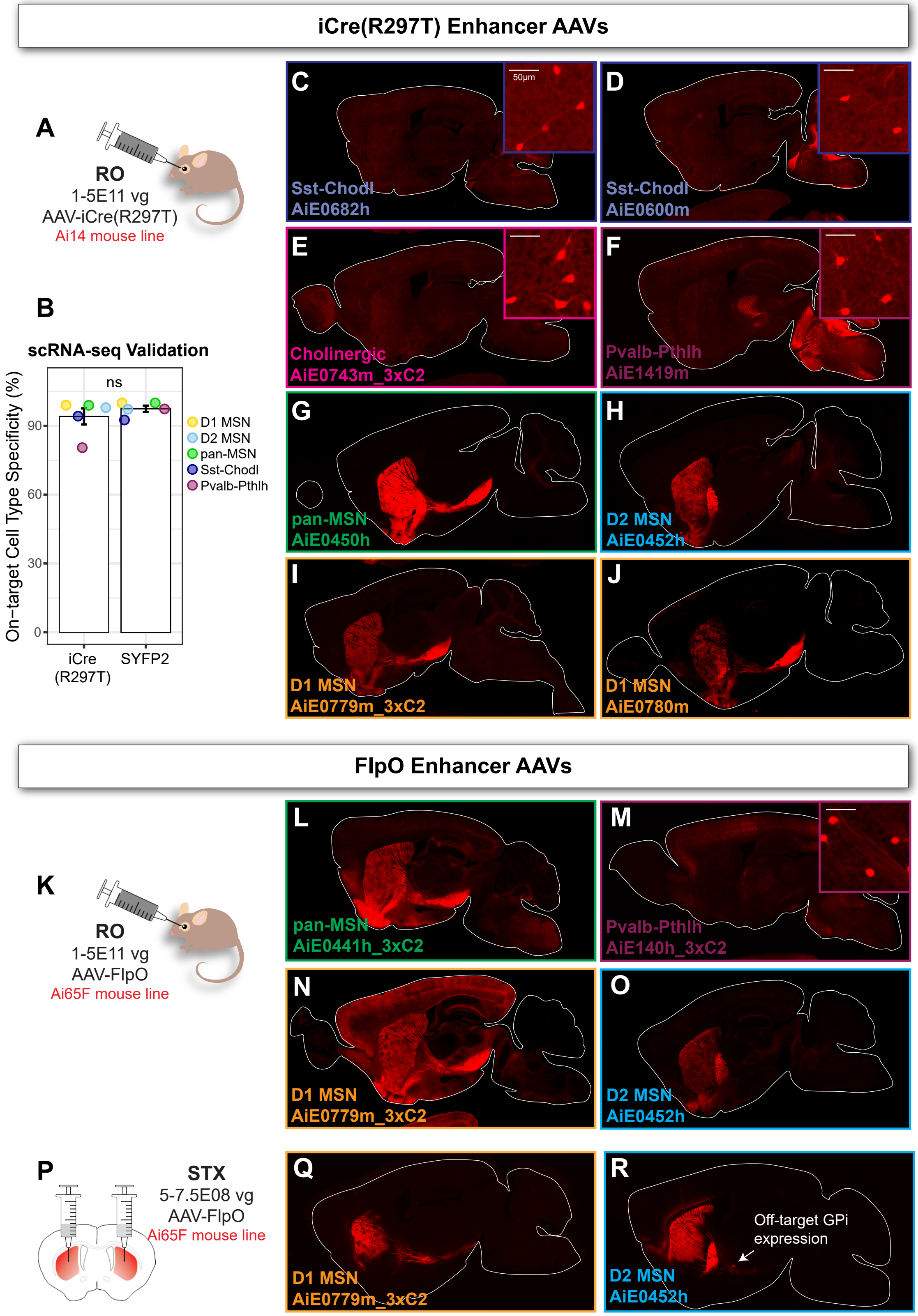


**Figure S7**: Striatal enhancer AAVs permit cell type-specific iCre or FlpO recombination.
A-J) Enhancer AAVs driving iCre(297T) injected RO into the Ai14 Cre-dependent tdTomato reporter mouse line.
A) Schematic of experiment. B) SSv4 scRNAseq validation of sorted cells showing no difference in cell type specificity of iCre(297T) vectors compared to matched SYFP2 reporter vector (Wilcoxon rank sum test, p=0.6752).
C-J) Example images of cell type specific tdTomato expression following enhancer driven Cre recombination.
K-R) Enhancer AAVs diriving FlpO injected into Ai65 reporter mouse line. K) Schematic of experiment for L-O.
L-O) Example images of cell type specific tdTomato expression following enhancer driven FlpO recombination.
P) Schematic of experiment for panels Q and R. Q-R) Example images of cell type specific tdTomato expression following enhancer driven FlpO recombination.

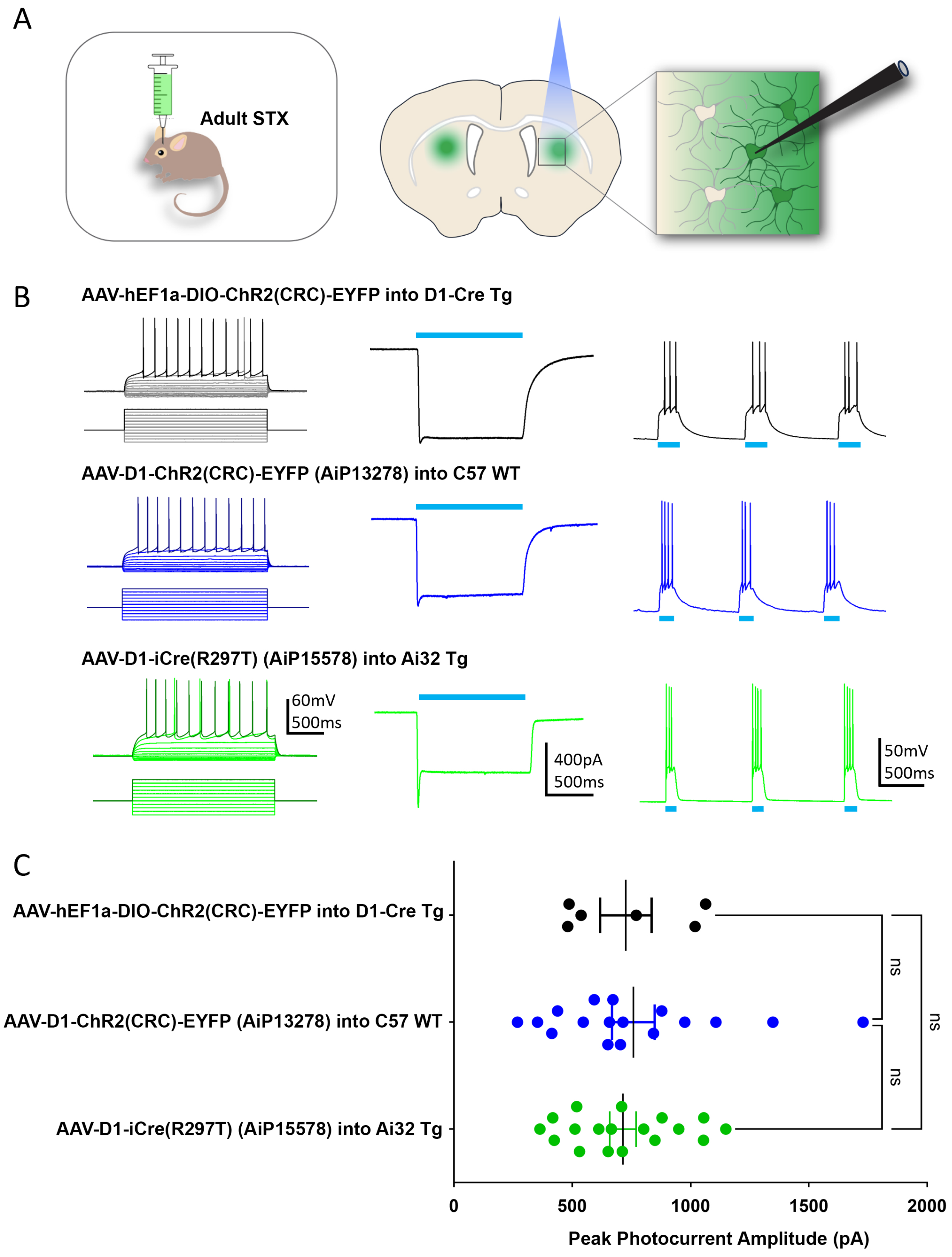


**Figure S8**: Comparison of ChR2 photocurrent amplitudes in D1 MSNs for different viral transgene expression strategies. A) Diagram of adult mouse stereotactic injection of AAV vectors into mouse brain (left) and acute brain slice recording set up with blue light stimulation applied to dorsal striatum region. B) Representative example whole cell current clamp recordings of ChR2 expressing D1 MSNs in response to a series of 1 second current injection steps, 50pA increments (left). Example whole cell voltage clamp recordings showing photocurrents evoked by 1 second blue light stimulation (middle). Example current clamp recordings showing multiple bouts of blue light-evoked action potential firing from the resting membrane potential. C) Summary data of peak photocurrent amplitude measurements for the different conditions as indicated. AAV-EF1a-DIO-ChR2(CRC)-EYFP virus injected into D1-Cre transgenic mice (n=6 cells; 1E+9vg dose), AAV-D1-ChR2(CRC)-EYFP virus AiP13278 injected into C57 wild-type mice (n=17 cells; 1E+9vg dose), and AAV-D1-Cre(R297T) virus AiP15578 injected into Ai32 Cre-dependent ChR2(H134R)-EYFP transgenic mice (n=18 cells; 2.5-5.0E+8vg dose). All viruses were packaged as AAV serotype PHP.eB. A one-way ANOVA revealed no significant difference between the three groups for mean photocurrent amplitudes ($p=0.9$). Tukey's multiple comparison test revealed no significant difference for any pairwise comparison between groups ($p \geq 0.9$ for all comparisons).

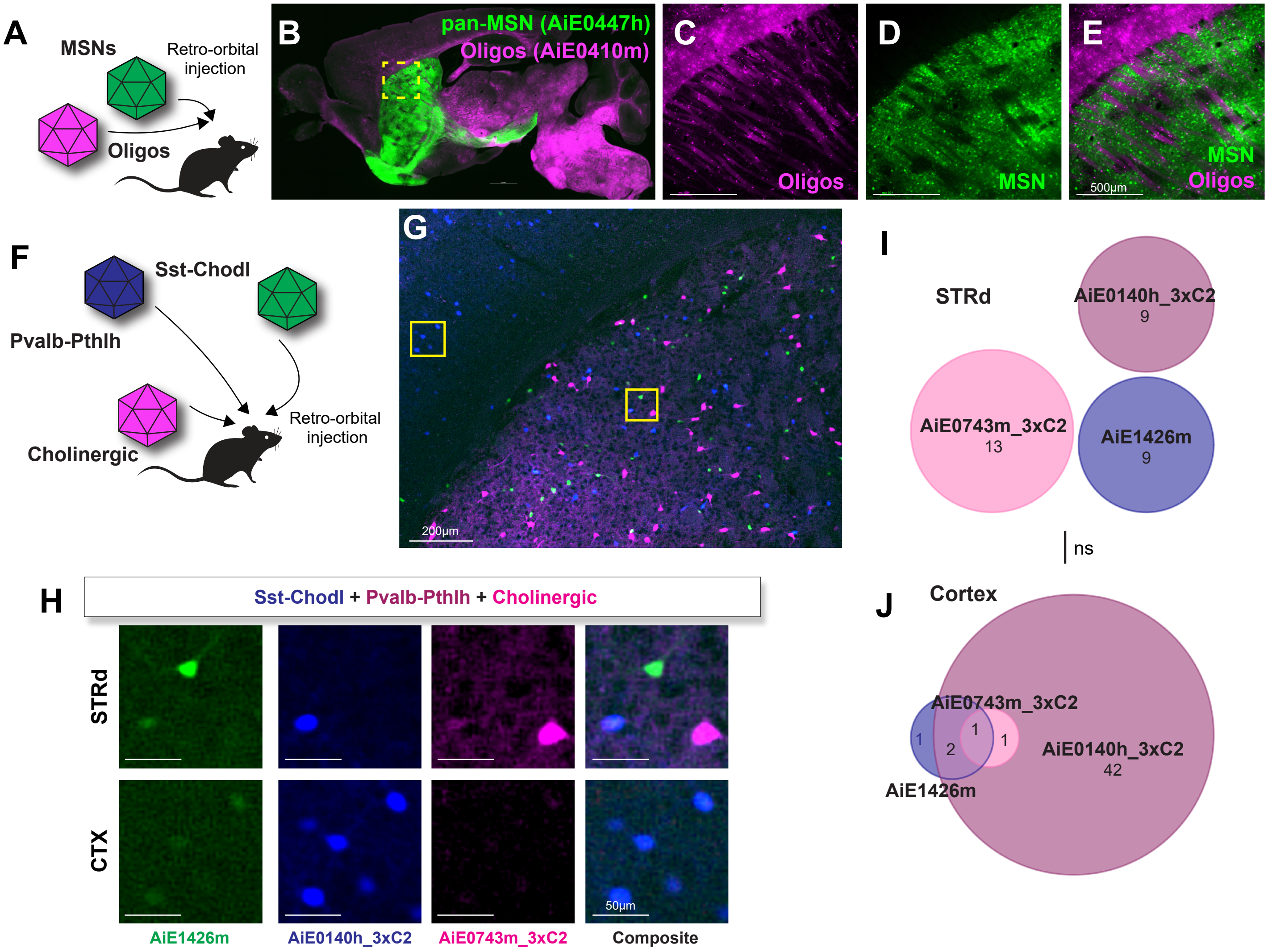


**Figure S9**: Successful multiplexing using different combinations of enhancer AAVs.
A-E) Pooling two enhancer AAVs for labeling pan-MSN (AiE0447h driving SYFP2) and oligodendrocytes (AiE0410m driving mTFP1). A) Diagram of the experimental paradigm. Enhancer AAVs were packaged separately using PHP.eB capsid and then combined for RO injection into an adult mouse (Oligo enhancer AiE0410m driving mTFP1, 7.5E+11vg; pan-MSN enhancer AiE0447h driving SYFP2, 5E+11vg). B) Sagittal section of SYFP2 and mTFP1 native fluorescence. C-E) Zoomed in views of dorsal striatum showing mutual exclusivity of enhancer labeling. F-J) Pooling three enhancer AAVs for labeling STR interneurons (Sst-Chodl enhancer AiE1426m driving SYFP2, 5E+11vg; cholinergic enhancer AiE0743m_3xC2 driving tdTomato 5E+11vg; and Pvalb-Pthlh enhancer AiE0140h_3xC2 driving mTFP1, 2E+11vg). F) Diagram of the experimental paradigm. Enhancer AAVs were packaged separately using PHP.eB capsid and combined for RO injection into an adult mouse. G) Native fluorescence from triple injection in cortex (CTX) and dorsal striatum (STRd). H) Zoomed in views from all three channels in STRd and CTX. I-J) Quantification of fluorescently labeled cells. Number of cells counted are displayed underneath enhancer label. I) No overlap between enhancers was observed in STRd ROI. J) Most cells in cortex ROI were mTFP1 positive (Pvalb interneurons, AiE0140h_3xC2), with only a few cells expressing off-target SYFP2 and tdTomato. To evaluate enhancer expression overlap in cortex vs STRd (n=1 mouse and ROI per region), Barnard's test was used. No significant differences were observed (p=0.24, two-sided).

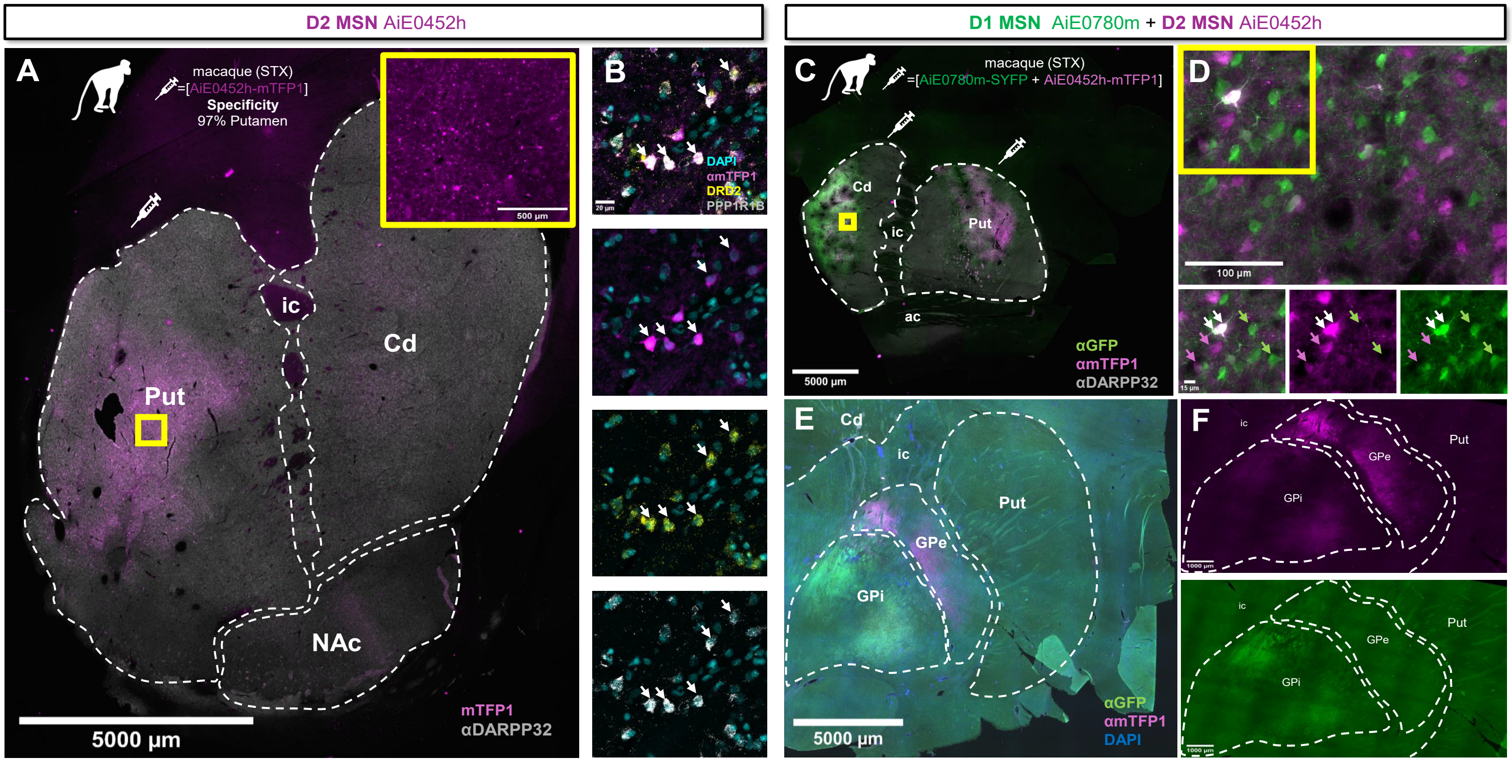


**Figure S10**: Single and duplex enhancer AAV injections label specific MSN populations in macaque striatum A-B) Stereotaxic injection of AiE0452h driving mTFP1 in macaque putamen. The total viral dose was 1E+11vg into putamen. A) Coronal section of the striatum region with mTFP1 (magenta) and αDARPP32 (grey). Inset of putamen shows native mTFP1 signal (magenta). B) RNAscopeV2 co-detection validation of AiE0452h with PPP1R1B and DRD2 probes. Cyan=DAPI, magenta=αmTFP1, yellow=αDRD2, and grey=αPPP1R1B. Arrows point to examples of mTFP1+/DRD2+/PPP1R1B+ cells. C-F) Duplex stereotaxic injection of mixture of AiE0780m driving SYFP2 and AiE0452h driving mTFP1 in macaque caudate and putamen. The total viral dose was 1E+11vg into caudate and 1E+11vg into putamen regions (5E+10vg for each virus per site). C) Coronal section of the striatum with αDARPP32 (grey), αGFP (green), and αmTFP1 (magenta). D) Top: Zoomed in view of cell labeling from injection into the caudate region. Bottom: Views of all channels separately from top inset. Arrows point to representative cells with distinct labeling of SYFP2+ (green arrows), mTFP1+ (purple arrows), and co-labeled SYFP2+/mTFP1+ (white arrows) cells. E-F) Coronal sections posterior to the injection site showing axon projections terminating in the GPi or GPe indicative of D1 (SYFP2+) and D2 (mTFP1+) MSNs of the direct and indirect pathways, respectively. All injections were performed using a BrainSight (Rogue Research) surgical robot. Abbreviations: Cd, caudate; Pu, putamen; NAc, nucleus accumbens; ic, internal capsule; GPe, globus pallidus external segment; GPi, globus pallidus internal segment.
